# Distributed state-dependent neural ensembles across sleep stages

**DOI:** 10.64898/2026.08.13.744563

**Authors:** Mirna Merkler, Alba Pascual Clavel, Shuzo Sakata

**Affiliations:** Strathclyde Institute of Pharmacy and Biomedical Sciences, University of Strathclyde 161 Cathedral Street, Glasgow G4 0RE, UK

## Abstract

Sleep architecture is organized by neural ensembles operating across multiple timescales, yet the organizing principles remain unclear. By monitoring neural populations across >40 brain regions in mice, we reveal a distributed sleep code spanning multiple temporal scales. On a slow (minutes-to-seconds) timescale, vigilance state reorganized brain-wide firing and was decodable from every region, with REM sleep emerging as a globally activated state. State transitions followed low-dimensional trajectories, with cortical and subcortical ensembles evolving in antiphase at wake–NREM boundaries but in register during transitions into REM sleep. On a fast (seconds-to-milliseconds) timescale, NREM slow/delta oscillations served as a global rhythm while hierarchically nesting spindles, sharp-wave ripples, and pontine (P) waves, whereas REM theta–P-wave coupling coordinated firing across regions. Regional sleep-related activities covaried with neuromodulatory innervation, while infraslow, history-dependent firing dynamics tracked NREM–REM cycles. Together, these findings provide a cell-resolved atlas of distributed neural population dynamics across sleep stages.

## Introduction

Sleep is conserved across the animal kingdom and is essential for cognition, memory consolidation, and homeostatic functions ^1–3^; correspondingly, disrupted and fragmented sleep accompanies a wide range of neurological and psychiatric disorders ^4–7^. While mammalian sleep is classified into two macroscopic stages, non-rapid eye movement (NREM) and rapid eye movement (REM) sleep ^8–11^, our understanding of how sleep is organized at the single-neuron level remains fragmented.

Sleep is structured by neural activity operating over multiple timescales. On the longer, circadian timescale, the two-process model frames sleep as the interaction of a homeostatic “Process S” and a circadian “Process C” ^12,13^. Embedded within this cycle is the ultradian rhythm, which governs the continuous shift between NREM and REM sleep ^9,14–18^. The regulatory mechanisms of the ultradian rhythm remain incompletely understood and have been attributed to a dedicated central pattern generator ^19^, to mutually inhibitory REM-on and REM-off populations ^10,16,20^, and to the homeostatic accumulation and discharge of REM propensity, an “hourglass”, in which REM pressure builds during NREM and is released during REM sleep ^18,21–26^.

Superimposed on this architecture, sleep is further structured by subsecond-to-second oscillations and events. The cardinal NREM signature is the <1 Hz cortical slow oscillation ^27^, which propagates as a travelling wave ^28,29^. Slow oscillations, sleep spindles, and hippocampal sharp-wave ripples (SWRs) are often coupled, and their coordination has been implicated in memory consolidation ^30–34^. A slower infraslow rhythm, on the order of tens of seconds, paces the alternation between states by gating the NREM-to-REM transition ^15,35,36^.

REM sleep, by contrast, is organized by hippocampal theta rhythms ^37,38^ together with ponto-geniculo-occipital waves (termed P-waves in rodents) that accompany REM and are phase-coupled to theta rhythms ^39–43^. However, P-waves are not strictly stage-specific; they also occur during NREM sleep, where they couple with SWRs in rodents ^42,44^ and macaques^45^.

Resolving how distributed neural ensembles coordinate across these events and state transitions requires simultaneous single-unit recordings across multiple brain regions. High-density silicon probes now make this approach feasible ^46,47^, reopening fundamental questions about functional architecture across different brain states. Although classical frameworks view the brain as a mosaic of functionally segregated regions, recent brain-wide mapping has challenged this localized model ^48–50^. An emerging view is that behavioral and cognitive variables are represented in a distributed but non-uniform fashion across the brain, concentrating in specific circuits rather than a single region, while retaining low-dimensional structure ^51,52^.

However, this network-level organization has been investigated primarily in the awake brain. Whether sleep stages, transitions, and their ultradian homeostatic processes rely on dedicated circuits or are sustained by a distributed, brainwide code remains unknown. Here, we address this gap by combining Neuropixels recordings in mice, tracking single-unit activity across more than 40 regions spanning from the isocortex to the pons. We specifically examine: (1) how population firing distinguishes vigilance states across the brain; (2) the coordination of cortical and subcortical ensembles during state transitions; (3) the multi-regional signatures of sleep oscillations and events; (4) the relationship between neuromodulatory anatomy and this functional organization; and (5) how distributed neural ensembles represent the homeostatic regulation of REM sleep.

## Results

To obtain single-unit activity from multiple brain regions during sleep-wake cycles, we performed acute Neuropixels recordings in head-fixed mice (n = 22 mice, 62 probe penetrations) with simultaneous frontal electroencephalogram (EEG), nuchal electromyogram (EMG) and pupillometry (**Fig. 1a**). Mice cycled spontaneously through wakefulness, NREM sleep and REM sleep during recording sessions lasting ∼4 hours. Vigilance states were scored in 4-s epochs based on the EEG/EMG signals; the overall sleep architecture is summarized in **Extended Data Figure 1**. During a recording, we typically observed state-dependent population firing across regions (**Fig. 1b**). Following the conclusion of recording sessions, probe trajectories were histologically reconstructed (**Fig. 1c**) and registered to the Allen Common Coordinate Framework (CCF) ^53^ (**Extended Data Fig. 2**).

**Figure 1.**
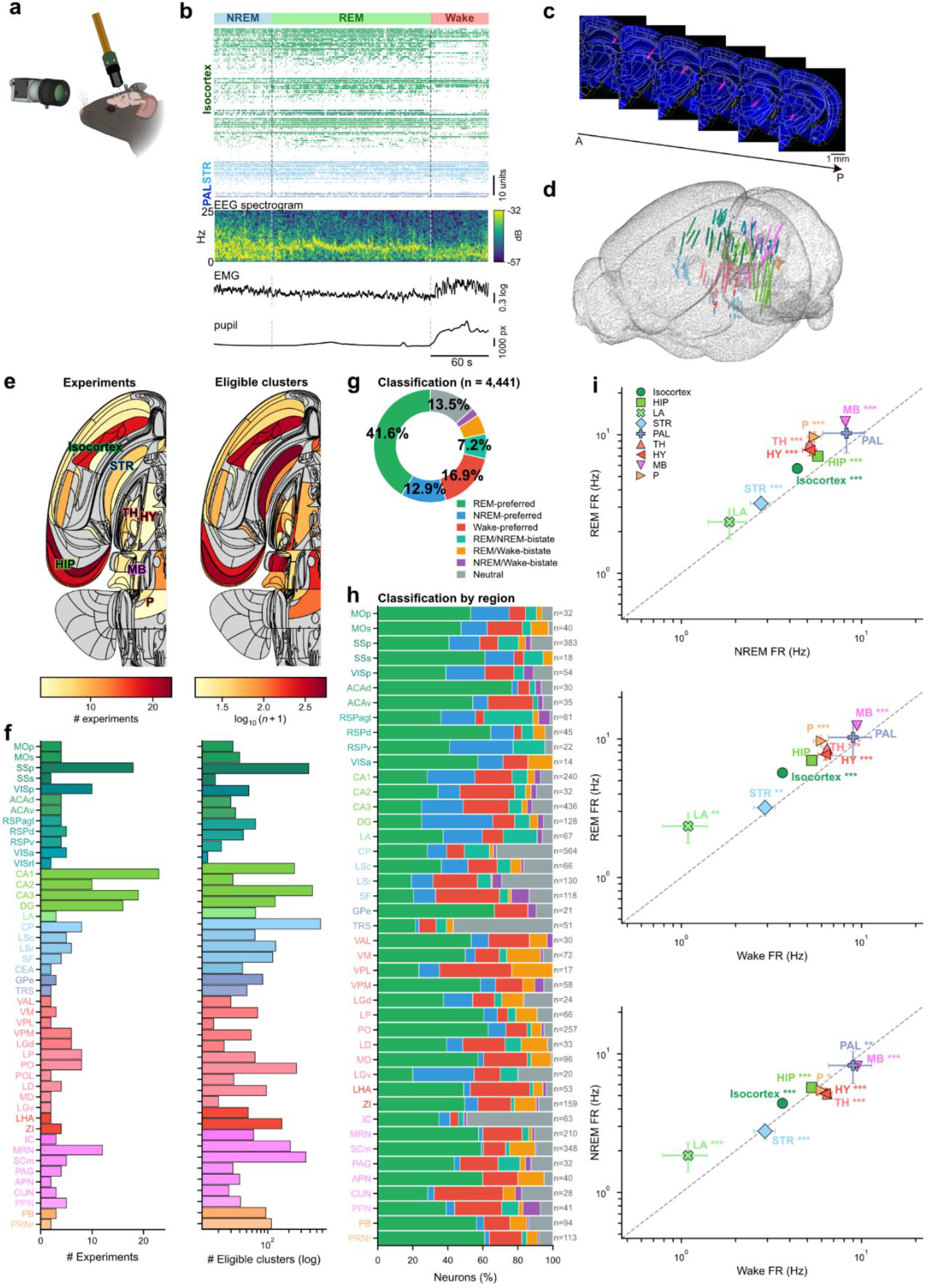
Database and vigilance state-dependent firing across brain regions. (**a**) Experimental schematic. (**b**) Example recording with three vigilance states, spike raster, cortical EEG spectrogram, EMG power and pupil diameter profiles. (**c**) Representative coronal histological sections showing the probe track. (**d**) Three-dimensional mouse brain rendering of all 4,555 localized single units colored by high-level structure. (**e**) Swanson dorsal-cortical and sub-cortical flatmaps showing (left) number of recordings and (right) number of eligible single units per region (log-scaled colour). (**f**) Per-structure summary bars showing number of single units (left) and number of recordings (right) for the nine well-sampled high-level structures. (**g**) Cohort-wide single-unit classification (n = 4,441; permutation test, 10,000 permutations, Benjamini-Hochberg False Discovery Rate (BH-FDR) α = 0.05, bistate |log_2_FC| < 0.5). Donut: proportion in each of seven categories. Stacked bar: same data as absolute counts. (**h**) Stacked bar plot of the same classification, broken down by region. (**i**) Per-structure mean firing rate across state pairs (top: REM vs NREM; middle: REM vs Wake; bottom: NREM vs Wake) on log–log axes; markers are mean ± s.e.m. per high-level structure. Asterisks denote paired Wilcoxon signed-rank tests across the nine high-level structures, the unit of inference for this panel, with BH-FDR across structures (*p < 0.05, **p < 0.01, ***p < 0.001).

Following spike sorting and quality control (QC) (**Extended Data Fig. 3**), 4,555 single-units passed QC thresholds and were assigned to CCF regions. These units spanned nine high-level structures: isocortex (n = 740 units across 12 regions), hippocampus (HIP, n = 836; 4 regions), lateral amygdalar nucleus (LA, n = 67; 1 region), striatum (STR, n = 922; 5 regions), pallidum (PAL, n = 136; 2 regions), thalamus (TH, n = 673; 11 regions), hypothalamus (HY, n = 212; 2 regions), midbrain (MB, n = 762; 7 regions) and pons (P, n = 207; 2 regions). Those sampled high-level structures and individual brain regions are summarized in three-dimensional CCF reconstructions (**Fig. 1d**), a Swanson flat-map ^54^ (**Fig. 1e**; see also **Supplementary File**) and per-structure yield summaries (**Fig. 1f**).

### REM sleep as a global activation state

To compare gross activity across states and regions, we statistically assessed state-dependent changes in firing rates using a permutation framework (see Methods). Among the neurons that met our threshold of at least five 4-s epochs (i.e., 20 s) per state (n = 4,441 neurons from 59 recordings), 86.5% showed significant state modulation. Specifically, 71.4% were classified into a single preferred state, while 15.1% fell into a bi-state category (**Fig. 1g**). REM-preferred neurons constituted the largest functional class (41.6%), followed by Wake-preferred (16.9%) and NREM-preferred (12.9%) neurons; only 13.5% remained neutral, showing no significant modulation.

Although these functional categories were heterogeneously distributed, REM-preferred neurons predominated in most recorded regions (**Fig. 1h**). This global REM activation became more pronounced when we compared mean firing rates across states (**Fig. 1i** and **Extended Data Fig. 4**). REM exceeded both NREM and wake across the nine high-level structures (W = 0, p = 3.9 × 10^-3^ for both, two-sided Wilcoxon signed-rank), whereas NREM and wake did not differ (W = 17, p = 0.57). Thus, despite diverse functional cell types within regions, REM sleep represents a global activation state.

### Decoding of vigilance states by distributed activity

Next, we asked whether vigilance states could be inferred from population spike counts alone, and how this information is distributed across the brain. We trained Random Forest classifiers taking two complementary approaches (**Fig. 2a**): (1) an all-neuron decoder using all simultaneously recorded neurons to classify each 4-s epoch, and (2) a region-restricted decoder using only neurons within a single brain region. For the all-neuron decoder, we also quantified individual neural contributions by computing Gini feature importance.

**Figure 2.**
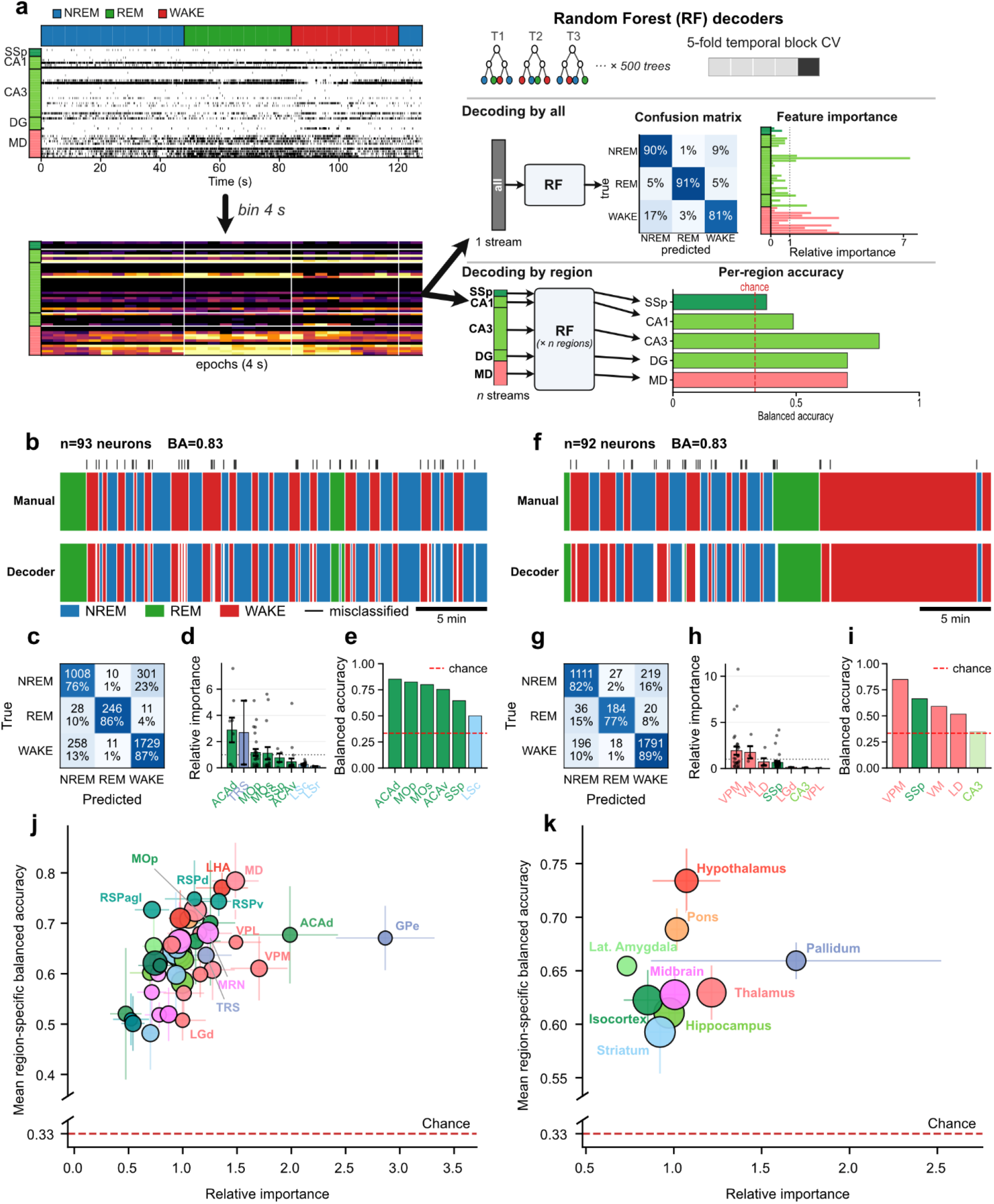
Vigilance state decoding. (**a**) Decoding pipeline. Top-left: example spike raster with the scored vigilance state shown above. Bottom-left: same data binned at 4 s and z-scored to produce the decoder input matrix (neurons × epochs); Right: Random Forest (RF) classifier (500 trees) with 5-fold temporal-block cross-validation, yielding a confusion matrix and per-unit relative Gini feature importance for an all-neuron decoder, and the balanced accuracy (BA) for a region-specific decoder. (**b–i**) Decoding outputs from two example recordings. (**b, f**) Manually curated (top) and decoded (bottom) state hypnograms; tick marks flag misclassified epochs. (**c, g**) Row-normalized confusion matrix. (**d, h**) Mean relative RF feature importance grouped by source region; dotted line, 1.0 = a neuron of average importance in that recording. (**e, i**) Region-restricted BA for each contributing region (dashed line: chance, 0.33). (**j**) Region-level relationship between feature importance and standalone decoding. Each marker is one Allen CCF region, aggregated across recordings; relative Gini importance and standalone accuracy were positively related (Spearman’s ρ = 0.564, p = 8.1 x 10^-5^, n = 43 regions). Marker area encodes the number of contributing units, and marker edge color whether that region’s own decoder exceeded chance in at least half of the recordings sampling it (black) or fewer (white); error bars, s.e.m. (**k**) Same plot at the level of the 9 high-level Allen structures (markers are neuron-weighted means across all constituent regions and recordings; bubble area encodes the number of contributing units). At this coarser level, the same relationship was not resolved (Spearman’s ρ = 0.533, p = 0.14, n = 9 structures).

We first analyzed two representative recordings - one heavily sampling cortical neurons (**Figs. 2b-e**) and the other subcortical neurons (**Figs. 2f-i**). In both instances, the all-neuron decoder achieved high overall accuracy (83%; **Figs. 2b,f**) with robust recall across all states (**Figs. 2c,g**). Mean relative feature importance was variable across the sampled regions (**Figs. 2d,h**). Region-restricted decoders also performed consistently above chance across most individual regions (**Figs. 2e,i**), suggesting that robust state information is preserved locally.

Applying this approach across the entire cohort confirmed the widespread distribution of sleep-state information. While decoding accuracy increased with the number of simultaneously recorded neurons and brain regions (**Extended Data Fig. 5**), all-neuron decoders maintained high accuracy across all sessions (72.5 ± 1.3%, range: 51.5 - 87.2%). This positive neuron-count relationship held within the isocortex, HIP, TH, HY, and MB (**Extended Data Fig. 5**). The mean of relative importance across regions significantly correlated with the mean region-specific balanced accuracy (Spearman ρ = 0.564, p = 8.1 x 10^-5^, n = 43; **Fig. 2j**). While the HY and Pons tended to show higher balanced accuracy (**Fig. 2k**), across 181 region-restricted decoders (individual regions × recordings), 97.2% performed significantly above chance (permutation test, *p* < 0.05), and every tested region was significant (Fisher’s combined test), suggesting that vigilance state is represented by distributed neural population activity.

### Dissociation of cortico-subcortical populations during state transitions

We next examined how population activity reorganizes during state transitions. We aligned neuronal activity across all regions, along with simultaneous frontal EEG and EMG, to the boundaries of four canonical transitions: Wake → NREM, NREM → Wake, NREM → REM and REM → Wake (**Fig. 3**). Population EEG spectrograms confirmed the canonical spectral signatures of each transition (**Figs. 3a–d**). Z-scoring regional firing rates relative to a pre-transition baseline (−60 to −30 s) revealed widespread yet heterogeneous reorganization across regions (**Figs. 3e–h**).

**Figure 3.**
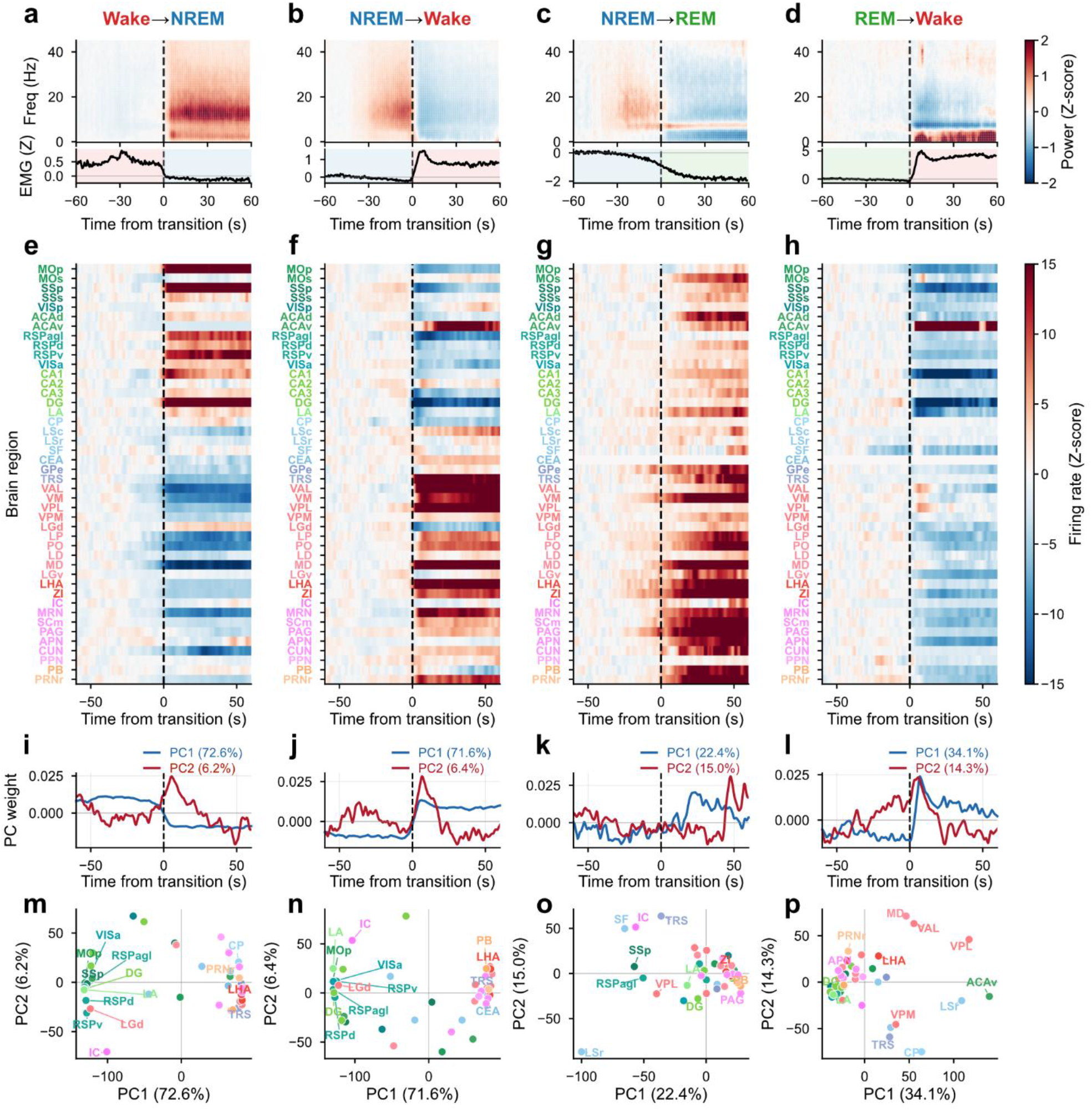
Neural population activity around the four sleep-state transitions. Columns, left to right: Wake → NREM, NREM → Wake, NREM → REM and REM → Wake. (**a–d**) Population EEG spectrogram (power z-scored to the −60 to −30 s baseline; diverging color map saturated at ± 2 z) with the mean nuchal EMG (z-scored) below; vertical dashed line marks the scored transition boundary (t = 0). (**e–h**) Firing-rate z-score heatmaps: each row is one region. The z-scored pure-state regional profile was color-coded (10-ms bins; NaN-aware Gaussian smoothing, FWHM 2 s; per-region z-score against the −60 to −30 s baseline; diverging map saturated at ± 15 z). (**i–l**) Temporal weight vectors of the first two PCs of the regional profiles. (**m–p**) Projection of every region onto PC1 (x) and PC2 (y) for the matching transition. Neurons with < 0.1 Hz in-window mean rate excluded; PCA computed independently per transition on 44 regions (Wake↔NREM) / 43 regions (REM transitions). n = 22 mice, 62 probe penetrations (60 with REM).

To extract the dominant temporal features underlying these regional dynamics, we applied principal component analysis (PCA) to the region × time matrix of these profiles, separately for each transition (**Fig. 3i–p**). At Wake → NREM and NREM → Wake transitions, population dynamics were low-dimensional, with the first principal component (PC1) explaining >70% of the variance. Along the PC1 axis, cerebral cortical structures (Isocortex, HIP, and LA) and the remaining subcortical regions segregated into opposite poles (**Figs. 3m,n**). This reciprocal relationship was consistent: during Wake → NREM transitions, cerebral cortical populations increased their firing rates while subcortical activity decreased, whereas the inverse pattern occurred during NREM → Wake transitions.

In contrast, REM-related transitions (NREM → REM and REM → Wake) exhibited qualitatively distinct dynamics: the leading components explained less variance (**Figs. 3k, l**), and the activity shifted uniformly in the same direction across most regions (**Figs. 3o,p**). Crucially, we observed distinct cortico-subcortical coordination: during NREM → REM transitions, subcortical regions exhibited larger changes in activity than cortical regions, whereas cortical regions showed greater modulation during REM → Wake transitions.

To visualize these region-specific trajectories, we re-embedded each firing-rate profile into a shared delay-embedded PC space to decompose the shared time course rather than the differences between regions (see Methods) (**Fig. 4 and Supplementary Video**). The resulting trajectories were low-dimensional, with PC1 alone capturing 96.8–99.3% of the variance across all four transitions (**Figs. 4a–d**). During NREM transitions (Wake → NREM and NREM → Wake), cortical and subcortical populations moved in opposite directions along the PC1 axis, with highly variable displacement magnitudes across regions (**Figs. 4a,b**). During the NREM → REM transition, most regions advanced in the same direction along PC1, led by large displacements in brainstem structures (**Fig. 4c**). Conversely, during the REM → Wake transition, most regions shifted in the opposite direction along PC1, dominated by large cortical displacement (**Fig. 4d**).

**Figure 4.**
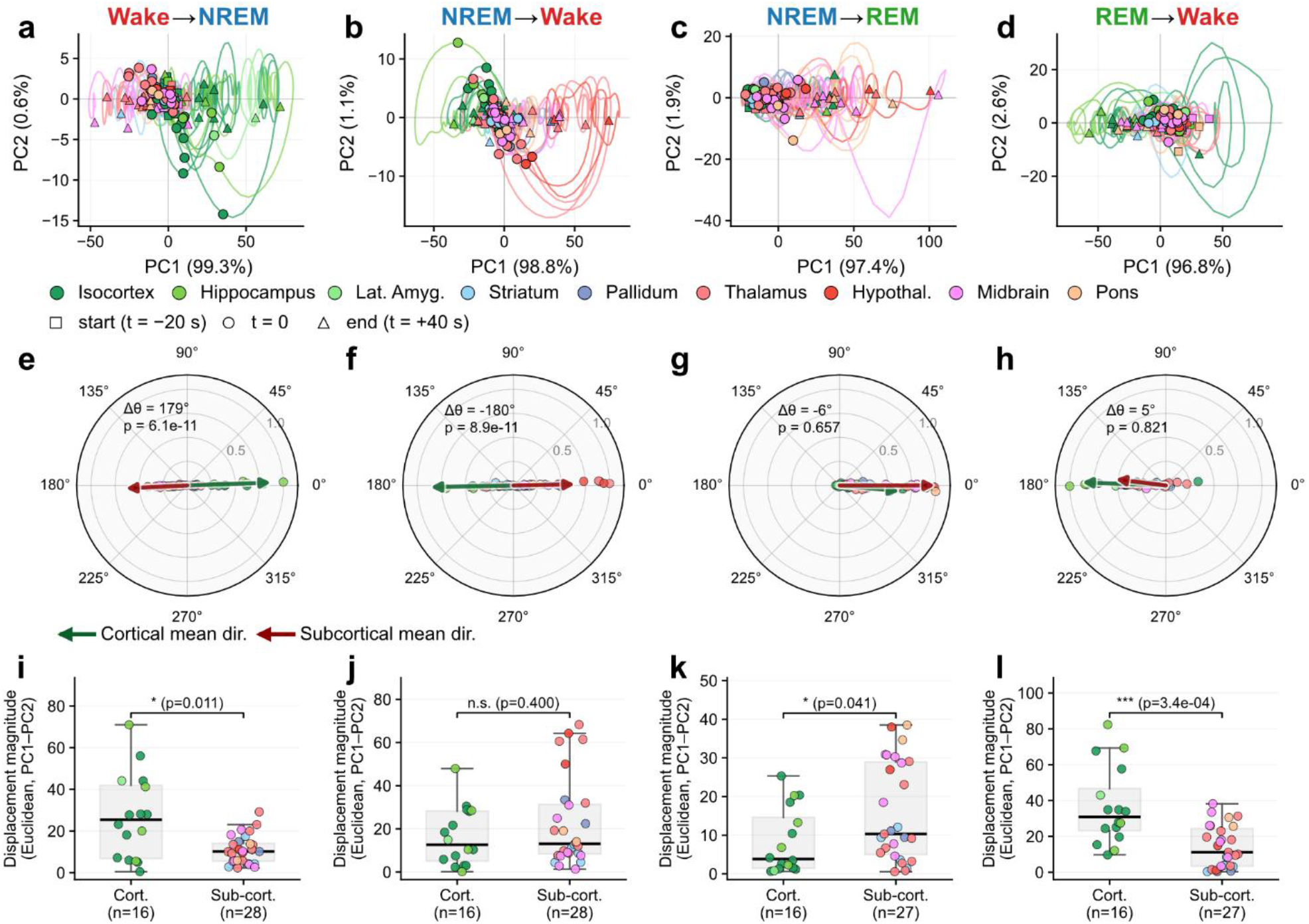
Cerebral cortical and subcortical population trajectories around state transitions. *Columns*, left to right: Wake → NREM, NREM → Wake, NREM → REM and REM → Wake. (**a–d**) Delay-embedded principal-component trajectories of each region’s peri-transition firing-rate profile (z-scored, FWHM 2 s, pure-state) in the PC1–PC2 plane; markers denote trajectory start (square, t = −20 s), the scored boundary (circle, t = 0) and end (triangle, t = +40 s); axis labels give the variance explained by each component; line and marker color encode the high-level structure. (**e–h**) Polar distribution of per-region displacement angles θ (direction of the pre- to post-transition centroid shift in the PC1–PC2 plane); the green and red arrows give the cerebral cortical and subcortical group mean directions (arrow length = mean resultant length, MRL; all distributions non-uniform, Rayleigh p < 0.002). The angular difference Δθ and its Watson–Williams p value are annotated. (**i–l**) Per-region displacement magnitude (Euclidean distance in the PC1–PC2 plane) for cortical versus subcortical regions; significance by two-sided Mann–Whitney U test (*p < 0.05, ***p < 0.001, n.s. not significant). Cortical group = isocortex, hippocampus and lateral amygdala (LA); subcortical group = striatum, pallidum, thalamus, hypothalamus, midbrain and pons. n = 22 mice, 62 probe penetrations (60 with REM).

We quantified these trajectories using displacement vectors from the pre-transition centroid (−20 to −5 s) to the post-transition centroid (+5 to +20 s) in the PC1–PC2 plane (**Figs. 4e-l**). For this analysis, regions were classified into a “cortical” group (isocortex, HIP and LA) and a “subcortical” group (STR, PAL, TH, HY, MB and P). During Wake → NREM and NREM → Wake transitions, the opposing trajectories of the two groups were significant (Wake → NREM: p = 6.1 × 10^-11^; NREM → Wake: p = 8.9 × 10^-11^, Watson-Williams test; **Figs. 4e,f,i,j**). Furthermore, cortical regions exhibited significantly larger displacement magnitudes during Wake → NREM transitions (p = 0.011, Mann-Whitney U-test). During REM-related transitions, the directions of the two groups were statistically indistinguishable (NREM → REM: p = 0.657; REM → Wake, p = 0.821). However, the displacement magnitude of the subcortical group was significantly larger during NREM → REM transitions (p = 0.041), but significantly smaller during REM → Wake transitions (p = 3.4 × 10^-4^) compared to the cortical group (**Figs. 4g,h,k,l**). Collectively, these findings demonstrate transition-dependent, region-specific population dynamics marked by a functional dissociation between cerebral cortical and subcortical networks.

### Multi-timescale, multi-regional coordination of NREM sleep-related neural events

Next, we examined how neural population activity is organized within each major sleep stage across multiple regions on a fast (seconds-to-milliseconds) timescale. During NREM sleep, we focused on the following oscillations and events: slow (<0.5 Hz) and delta (0.5-4 Hz) oscillations, sleep spindles, hippocampal SWRs and P-waves (**Fig. 5**). Although P-waves were traditionally recognized as a hallmark of REM sleep, they also occur during NREM sleep ^42,44,45^. We monitored P-waves via local field potentials using either a bipolar electrode or a Neuropixels probe (see Methods).

**Figure 5.**
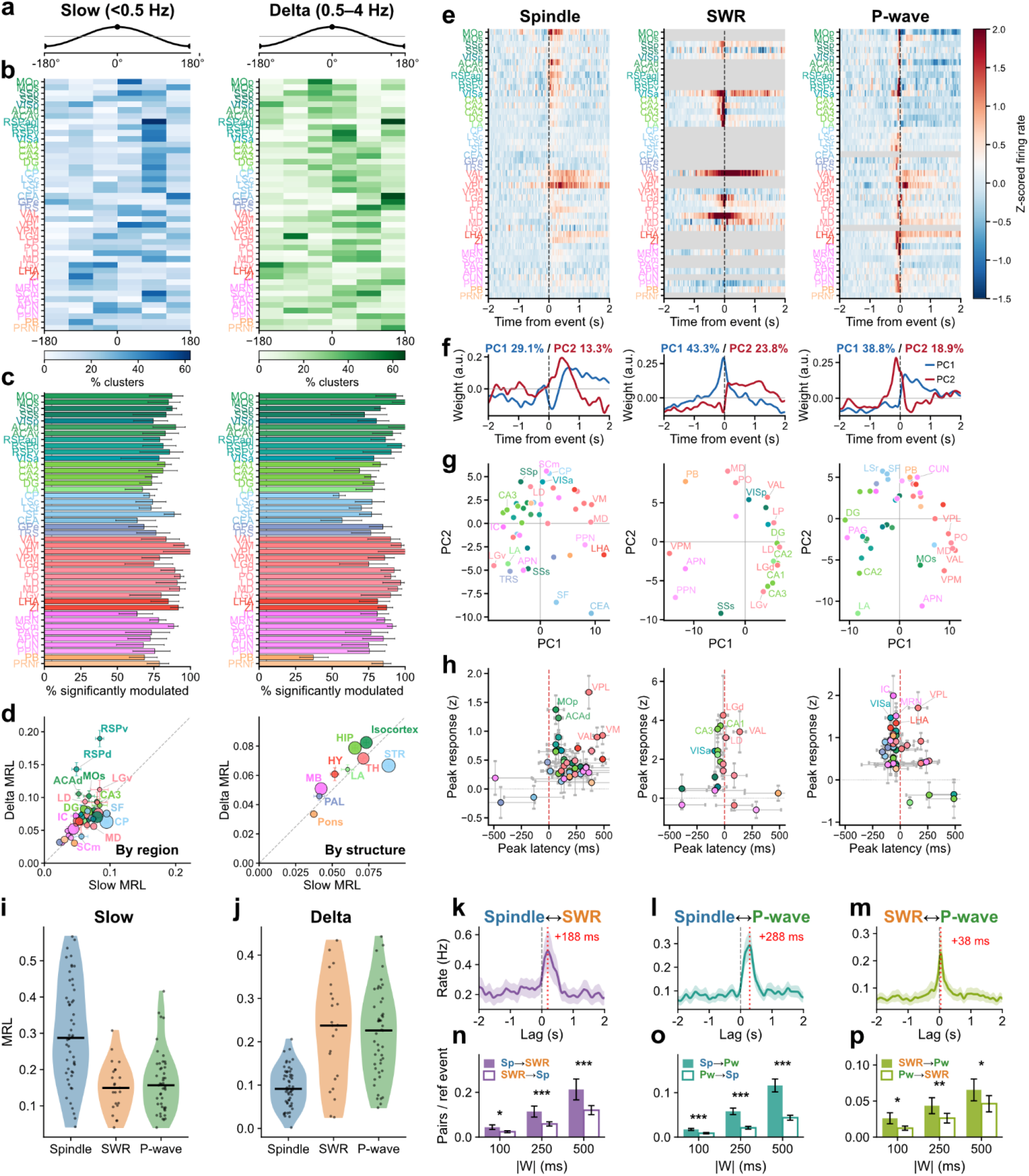
Multi-regional coordination of neural population activity across NREM sleep oscillations and events. (**a–d**) Single-unit coupling to the two NREM rhythms, slow (< 0.5 Hz) and delta (0.5–4 Hz) oscillations. (**a**) Reference cycle (peak/trough). (**b**) Phase-preference maps: for each region (rows; ≥ 5 units), the percentage of phase-modulated units preferring each 60° phase bin (slow, blues; delta, greens). (**c**) Percentage of units significantly phase-modulated per region (Rayleigh p < 0.05; horizontal lines, Wilson 95% CI). (d) Region-mean (left, n = 44) and structure-mean (right, n = 9) mean resultant length (MRL) for slow versus delta oscillations; point size scales with unit count. Paired region-level MRLs differed between the two rhythms (Wilcoxon signed-rank, W = 243, p = 0.003, n = 44 regions); the difference was not resolved across high-level structures (paired t-test, t = −0.71, p = 0.50, n = 9). (**e–h**) Event-triggered population dynamics for spindles, SWRs and P-waves (n = 4,555, 1,600 and 3,985 contributing units; ±2 s, 25-ms bins, z-scored). (**e**) Region-averaged firing-rate heatmaps (gray, region absent for that event). (**f**) PC1/PC2 temporal weight profiles from PCA. (**g**) PC1–PC2 region-loading scatter (points colored by Allen CCF scheme). (**h**) Per-region peak response amplitude versus peak latency. (i**,j**) Event-phase coupling strength (MRL per recording) of each event to the slow (**i**) and delta (**j**) oscillations; horizontal bars, medians; Kruskal–Wallis p = 6.8 x 10^-8^ (**i**), p = 1.3 x 10^-9^ (**j**). (**k–m**) Event–event cross-correlograms (mean ± s.e.m. across recordings; ±2 s, 25-ms bins; dashed line, t = 0; red, peak lag): spindle–SWR (**k**), spindle–P-wave (**l**), SWR–P-wave (**m**). (**n–p**) Directional cooccurrence: forward (filled) versus backward (open) partner events per reference event within ±100/250/500 ms windows (mean ± s.e.m.; paired Wilcoxon signed-rank per window; *p < 0.05, **p < 0.01, ***p < 0.001).

We first assessed how multi-regional single-unit spiking is entrained to the two major rhythms: slow and delta oscillations (**Figs. 5a-d**). Most neurons were significantly phase-locked to both rhythms (Rayleigh test, p < 0.05: 80.4% to the slow oscillation and 79.5% to the delta rhythm; **Figs. 5b,c**). To evaluate the magnitude of phase entrainment, we computed the mean resultant length (MRL) for each oscillation and compared population profiles across regions (**Fig. 5d**). MRL values were strongly correlated between the two oscillations (Spearman ρ = 0.64, p = 2.5 × 10^-6^, n = 44 regions; high-level structures: ρ = 0.88, p = 0.002, n = 9), although paired region-level MRLs still differed (Wilcoxon signed-rank, p = 0.003, n = 44). However, a subset of regions deviated from this trend: the midline cortical areas, including the retrosplenial area (RSP) and the anterior cingulate area (ACA), entrained more strongly to the delta rhythm, whereas subcortical regions like the caudoputamen (CP) and septofimbrial nucleus (SF) exhibited preferential entrainment to slow oscillations.

We next characterized multi-regional event-related activity across spindles, SWRs and P-waves (**Figs. 5e-h and Extended Data Figs. 6 and 7**). Spindles induced robust eventrelated activity in isocortical and TH regions alongside moderate modulations in HY and MB populations. SWR-related activity was evident across HIP, isocortical and TH regions. Notably, NREM P-wave-related activity was widespread.

To isolate the dominant modes of these event-related dynamics, we applied PCA (**Figs. 5f,g**). For spindles, PC2 captured a gradual increase in firing rates following event onset, heavily weighting regions in the isocortex, TH, HY and MB. For SWRs, the first two PCs captured distinct regional cascades: HIP populations exhibited tight phasic activity around the SWR peak (PC1), whereas the mediodorsal (MD) and posterior complex (PO) nuclei of TH showed delayed activation (PC2). For P-waves, MB and P regions showed early activation immediately preceding the P-wave trough (PC2), followed by delayed TH activation (PC1). Single-neuron peak response and latency analyses supported these patterns (**Fig. 5h**); TH populations exhibited later peak latencies during spindles (>250 ms post-onset), whereas strong responders to SWRs and P-waves displayed synchronized, narrow latency distributions.

Finally, we quantitatively assessed the temporal relationships across these discrete events (**Figs. 5i-p**). Within each band, the three event types differed in coupling strength; spindles were most strongly coupled to the slow rhythm (Kruskal–Wallis test across event types, p = 6.8 × 10^-^⁸), whereas SWRs and P-waves were biased toward the delta rhythm (p = 1.3 × 10^-9^; **Fig. 5i and j**). Cross-correlograms revealed non-random, asymmetric co-occurrence profiles with directional peak lags: SWRs followed spindles, P-waves followed spindles, and P-waves followed SWRs (**Fig. 5k–m**). Comparing forward versus backward co-occurrence probabilities within directional time windows confirmed this asymmetry (**Fig. 5n–p**), identifying a temporal sequence among the three distinct events (spindle → SWR → P-wave). These findings show that NREM sleep is characterized by structured, sequentially coordinated neural activity across the brain.

### Multi-regional coordination of REM sleep-related neural events

While REM sleep is characterized by theta rhythms and P-waves, how neural population activity aligns to these events has not been mapped at scale. We therefore quantified how single-unit spiking across the brain phase-locks to the theta rhythm and is modulated around P-waves (**Figs. 6a-g**). We found that significant theta phase-locking was widespread yet non-uniform (**Fig. 6a**); theta-modulated neurons were concentrated in the HIP, lateral septum (LS), TH, and isocortex (e.g., RSP), but were sparse in the hypothalamus (HY), brain-stem (MB and P), CP, and PAL. The magnitude of entrainment mirrored this anatomical distribution, peaking in the HIP and several TH nuclei (**Fig. 6b**).

**Figure 6.**
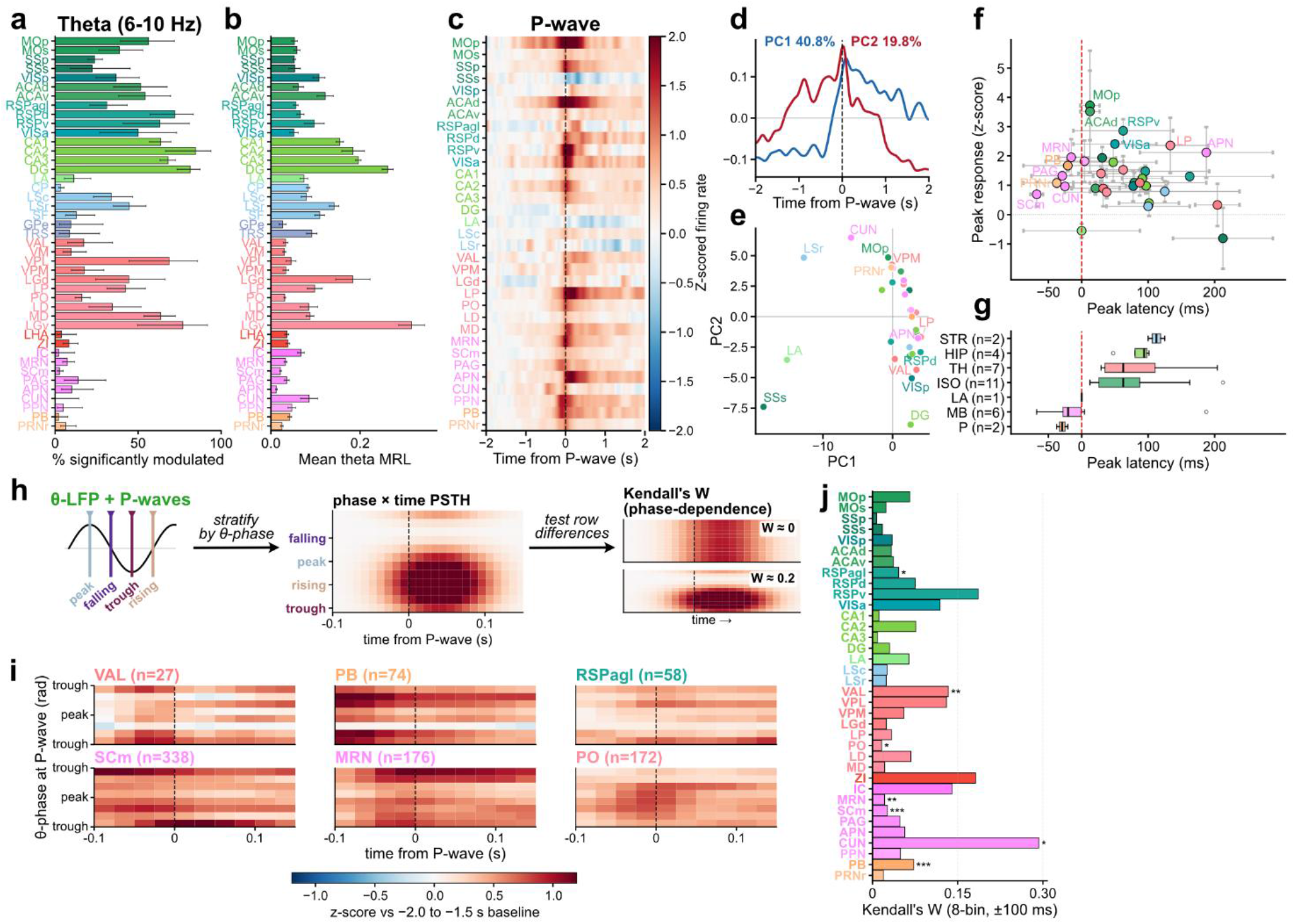
Multi-regional coordination of neural population activity across REM theta rhythms and P-waves. (**a**) Percentage of single units per region significantly phase-locked to theta during REM (Rayleigh test, Benjamini–Hochberg FDR within region; error bars, 95% confidence interval; n = 4,047 units, 43 regions). (**b**) Mean theta phase-locking strength (MRL) per region during REM (± s.e.m.). (**c**) Region-averaged perievent firing (z-scored to a −2 to −1.5 s baseline) aligned to REM P-waves (n = 8,961 P-waves; 33 regions; dashed line, P-wave trough timing). (**d**) Temporal weights of the first two PCs of the region-averaged peri-P-wave profiles. (**e**) Regions in the PC1–PC2 loading plane, colored by structure. (f) Per-region peak response amplitude versus peak latency (Spearman ρ = −0.18, p = 0.323, n = 33; dashed line, P-wave trough timing). **(g)** Peak latency by high-level structure (box, median and interquartile range; Kruskal–Wallis H = 12.38, p = 0.030, n = 33 regions in 7 structures). **(h)** Analysis schematic (P-waves stratified by theta phase; phase × time response matrix; Kendall’s W as a measure of theta-phase dependence). **(i)** Six example regions’ phase × time firing matrices (z-scored; ±100 ms window; 8 theta-phase bins). (**j**) Kendall’s W (8 phase bins, ±100 ms window) for each region tested. Kendall’s W and contributing unit count for the seven FDR-surviving regions: CUN 0.29 (n = 11), VAL 0.13 (n = 27), PB 0.07 (n = 74), RSPagl 0.05 (n = 58), SCm 0.03 (n = 338), MRN 0.02 (n = 176), PO 0.02 (n = 172). Asterisks: *FDR p < 0.05, **p < 0.01, ***p < 0.001, (Friedman test, BH-FDR across 36 regions).

Although previous studies showed that P-waves lead to multi-regional neural activation ^42,44,45,55–57^, P-wave-related spiking dynamics have not been mapped across the brain. We found that P-wave-related modulation is widespread (**Fig. 6c**). PCA revealed that PC1 captured activity aligned to the P-wave trough (time 0) and late-phase responses, whereas PC2 was biased toward early-phase activation (**Fig. 6d**). While most regions scored positively on the PC1 axis, reflecting global activation, a subset of MB and P regions scored positively on PC2, indicating early recruitment (**Fig. 6e**).

To chart this spatiotemporal cascade, we computed single-neuron peak responses and latencies (**Figs. 6f,g**). While response magnitude was uncorrelated with latency (Spearman ρ = −0.18, p = 0.323, n = 33 regions), peak latencies differed across high-level structures (Kruskal–Wallis H = 12.38, p = 0.030, 7 structures; **Fig. 6g**): P (median −29 ms) and MB (−20 ms) populations led the cascade, followed in order by TH, iso-cortex, HIP and STR. Thus, P-wave-associated firing sweeps from brainstem to forebrain over approximately 100–150 ms, roughly a theta cycle.

Consistent with previous reports ^42,58^, P-waves were theta-phase-locked, with theta power and frequency modulated around P-wave timing (**Extended Data Fig. 7**). However, brainstem neurons are strongly modulated by P-waves yet poorly entrained to theta (**Figs. 6a-c**), leaving it unclear whether theta-P-wave coupling reflects an active spike-level cross-regional code or merely an epiphenomenon at the local field potential level. We therefore tested region by region whether P-wave-related firing depended on the concurrent theta phase (**Fig. 6h**; see Methods). Seven of 36 regions showed significant phase-gating (BH-FDR; **Figs. 6i,j**), strongest by effect size in the cuneiform nucleus (CUN, Kendall’s W = 0.29), then the ventral anterior-lateral thalamus (VAL, W = 0.13), parabrachial nucleus (PB), RSP (RSPagl), superior colliculus (SCm), midbrain reticular nucleus (MRN) and posterior thalamic nucleus (PO). These results indicate that theta-P-wave coupling during REM sleep is distributed across high-level structures.

### Joint organization of neuromodulatory innervation and state-dependent activity

Although sleep-related activity is highly structured (**Figs. 5 and 6**), the organizational principles governing this distributed activity remain unclear. Because ascending neuromodulatory systems differentially innervate cortical and subcortical regions to modulate state-dependent firing, we hypothesized that regional neuromodulatory innervation density predicts these distinct electrophysiological signatures.

To test this, we used anterograde tracing data from the Allen Mouse Brain Connectivity Atlas ^59,60^ to quantify cell-type-specific projection volumes for four major ascending systems: dopaminergic (DA; ventral tegmental area and substantia nigra), serotonergic (5-HT; dorsal raphe), noradrenergic (NA; locus coeruleus) and cholinergic (ACh; pedunculopontine and laterodorsal tegmental nuclei) (**Fig. 7a** and **Extended Data Fig. 8**). We then correlated these volumes with six region-level metrics: NREM phase-locking MRL to slow and delta oscillations, and REM theta phase-locking MRL, P-wave firing modulation, P-wave response latency and theta–P-wave coupling (**Figs. 7b–g**).

**Figure 7.**
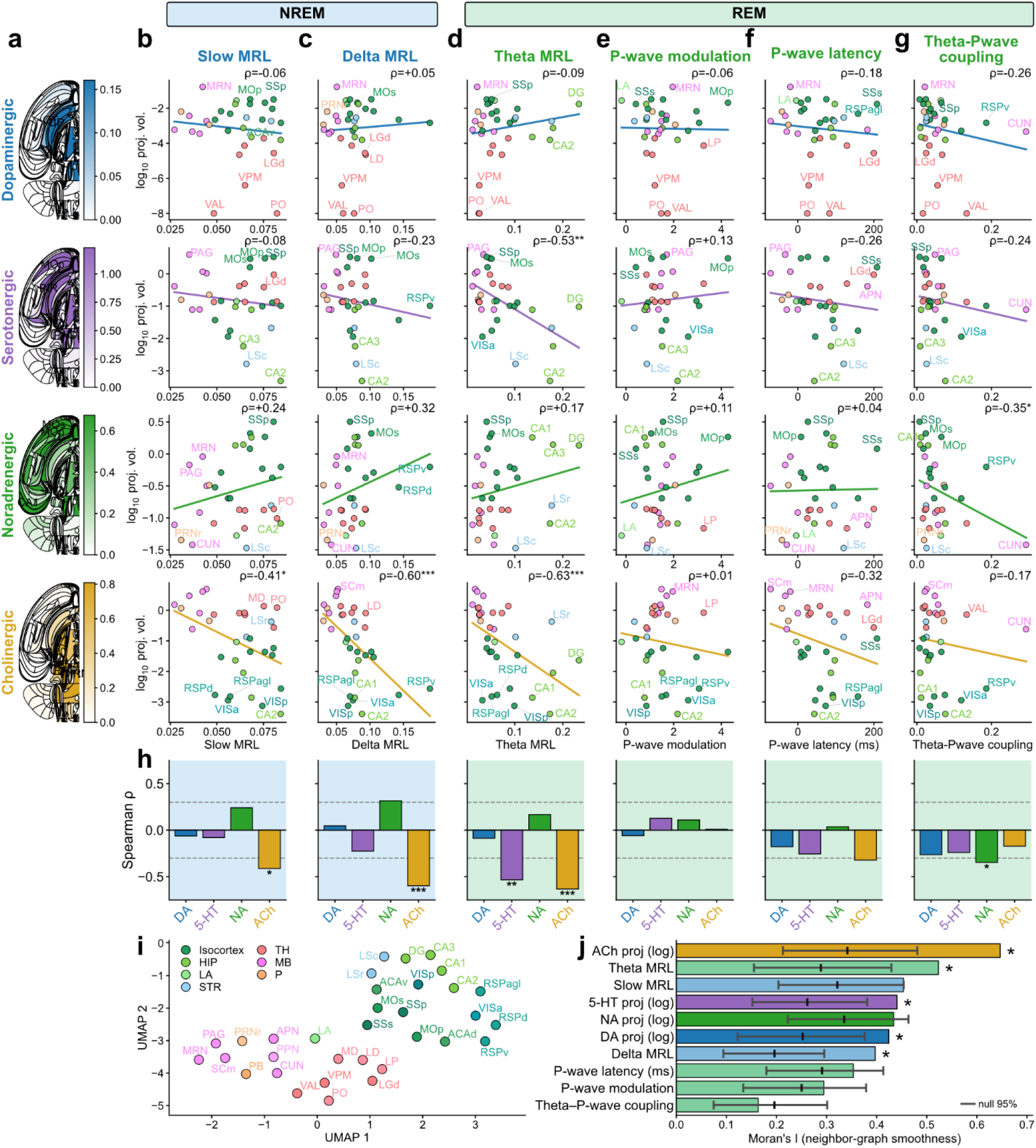
Joint analysis of ascending neuromodulatory innervation and electrophysiological sleep-related activity metrics. (**a**) Swanson flat-maps of cell-type-specific anterograde projection density for the four ascending neuromodulatory systems. top to bottom: dopaminergic, DA; serotonergic, 5-HT; noradrenergic, NA; cholinergic, ACh; maps are group means over n = 2 (DA), 2 (5-HT), 3 (NA) and 2 (ACh) Allen tracer experiments; injection cohort and whole-brain projection magnitudes in **Extended Data** Fig. 8. (**b–g**) For each neuromodulator system, per-region log10 normalized projection volume (y) versus a region-level sleep metric (x): (**b**) NREM slow phase-locking (MRL), (**c**) NREM delta phase-locking (MRL), (**d**) REM theta phase-locking (MRL), (**e**) REM P-wave firing modulation, (**f**) REM P-wave response latency and (**g**) REM theta–P-wave coupling (Kendall’s W, 8 phase bins, ±100 ms). Each point is one Allen CCF region colored by high-level structure; line, linear fit; top right, Spearman ρ (*p < 0.05, **p < 0.01, ***p < 0.001, uncorrected). (**h**) Spearman ρ for each system × metric (*p < 0.05, **p < 0.01, ***p < 0.001, uncorrected). Three of the 24 tests survive BH-FDR at q < 0.05: ACh × theta MRL, q = 0.002; ACh × delta MRL, q = 0.003; 5-HT × theta MRL, q = 0.011. (**i**) UMAP embedding of the 33 regions in the joint projection × sleep-phenotype feature space. (**j**) Contribution of each feature to the embedding structure, quantified as Moran’s I on the ten-dimensional fuzzy neighbor graph that UMAP constructs. Asterisks, the five features exceeding their own null at BH-FDR q < 0.05 under both that design and a hold-out design in which each feature is scored on a graph built from the other nine alone (ACh projection, q = 0.002; theta MRL, q = 0.007; 5-HT projection, q = 0.010; DA projection, q = 0.010; delta MRL, q = 0.002).

Neuromodulatory innervation correlated selectively with regional MRLs, most strongly mesopontine cholinergic input (**Figs. 7b-d,h**): regions heavily innervated by cholinergic axons showed markedly weaker phase-locking to delta and theta oscillations (q = 0.003 and 0.002) but not slow oscillations (q = 0.083; n = 33 regions). Serotonergic projections were similarly negatively correlated with theta phase-locking (q = 0.011), noradrenergic inputs were positive but not significant (p = 0.35), and dopaminergic innervation showed no MRL relationship (all p ≥ 0.64, n = 31 regions). No neuromodulatory system correlated with P-wave firing modulation, response latency or theta–P-wave coupling after correction (all q ≥ 0.08; **Figs. 7e–h**).

To explore how brain regions organize within this joint 10-dimensional feature space, we embedded the 33 eligible regions using Uniform Manifold Approximation and Projection (UMAP; **Fig. 7i**). The unsupervised embedding recovered macro-anatomy, segregating structures into two major clusters: a telencephalic cluster (isocortex, HIP, LS) and a subcortical cluster (TH, LA, MB, P). Regions of the same anatomical group (hippocampal subfields, thalamic nuclei) also clustered together.

We ranked these ten features by Moran’s I, which quantifies how smoothly each feature varies over the neighbor graph underlying the embedding (**Fig. 7j**; see Methods). Moran’s I was positive for all ten metrics, but since the graph is itself built from those same ten features, positivity is expected under the null rather than evidence of contribution. Bench-marked against a permutation null that rebuilds the graph from the permuted data (see Methods), five of the ten exceeded their own null: mesopontine cholinergic innervation ranked highest (Moran’s I = 0.65, null mean 0.34, q = 0.002), followed by theta MRL (0.52, 0.29, q = 0.007), serotonergic (0.44, 0.26, q = 0.010) and dopaminergic innervation (0.42, 0.25, q = 0.010), and delta MRL (0.40, 0.20, q = 0.002). None of the P-wave-related metrics exceeded its null. Overall, this joint analysis recovered mesoscopic anatomical identity across regions.

### Infraslow, history-dependent distributed code of ultradian cycles

While we have characterized highly organized neural ensembles within NREM and REM sleep, the mechanisms governing the alternation between them, known as the ultradian cycle, remain poorly understood. An influential framework is Benington and Heller’s ‘hourglass model’ ^18,21^. This model posits that REM sleep pressure builds during NREM sleep and discharges during REM sleep; consequently, the duration of a REM episode (REM_pre_) should predict the length of the subsequent NREM interval (|N|) required to accumulate pressure for the next REM bout ^21–26^.

To test this prediction in our dataset, we analyzed macroscopic sleep architecture across 158 cycles from 19 mice (**Figs. 8a,b**). **Figure 8a** highlights two representative cycles: a short REM episode (REM_pre_ = 28 s) was followed by a brief NREM interval, whereas a prolonged REM episode (REM_pre_ = 152 s) preceded a lengthy NREM interval. Cohort-wide, the total subsequent NREM duration (|N|) scaled positively with REM_pre_ duration (Spearman ρ = 0.53, p = 9.8 × 10^−13^; **Fig. 8b**). This relationship persisted within animals, across cycle positions and between session halves (**Extended Data Figs. 9a-c**), but was absent in the shortest REM_pre_ tertile (β = −0.38; **Extended Data Fig. 9d**).

**Figure 8.**
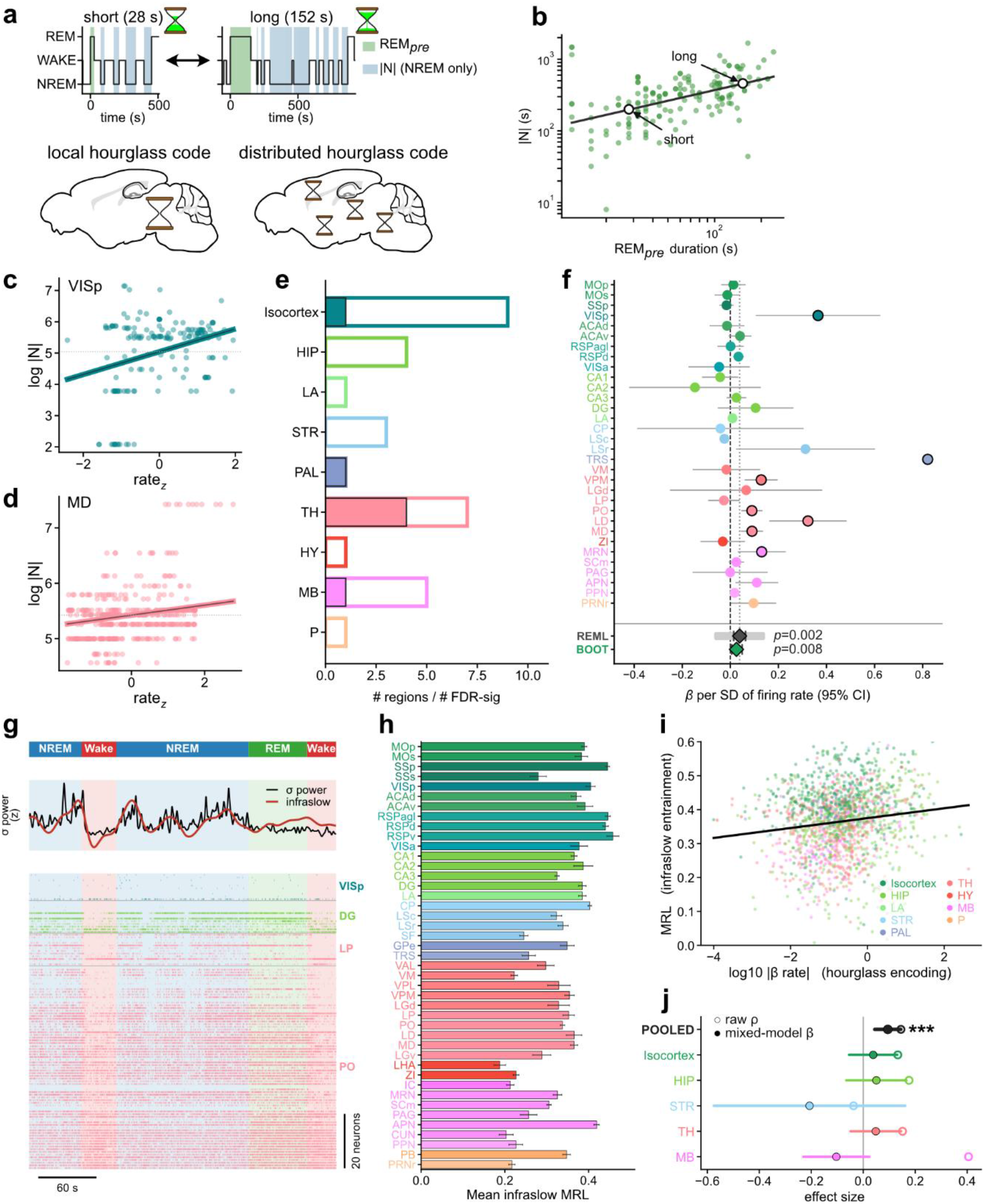
Infraslow, history-dependent distributed code for NREM-to-REM gating. (**a**) Top, example sleep cycles with a short (28 s) and a long (152 s) REM_pre_ episode (REM_pre_ shaded green; the NREM epochs summed into |N| shaded blue). Bottom, a schematic contrasting a local (single-hub) versus distributed (brain-wide) read-out of REM-homeostatic pressure. (**b**) Inter-REM NREM accumulation (|N|, total NREM time between the end of one REM episode and the start of the next) versus the duration of the preceding REM episode (REM_pre_); each point is one sleep cycle (n = 158 cycles, 19 mice; log–log axes; Spearman ρ = 0.53, p = 9.8 × 10^−13^; linear mixed-model slope β = 0.44, 95% CI [0.16, 0.72]). Arrows, examples in (**a**). (**c,d**) Example regions: log |N| versus within-unit z-scored REM_pre_ firing rate for primary visual cortex (VISp) (**c**) and mediodorsal thalamus (MD) (**d**); line, mixed-model fit. (**e**) Number of regions tested and significant, and number of significant units, grouped by high-level structure. (**f**) Per-region encoding slope (β per standard deviation of REM_pre_ rate) ± 95% CI, 32 regions; black rings, q < 0.05. REML, restricted maximum likelihood over the 28 regions with estimable SE (β = 0.039, 95% CI [0.014, 0.063]; pale bar, 95% prediction interval [−0.064, +0.141], p = 0.002). BOOT, the animal-cluster bootstrap (16 animals, effective 5.4; μ = +0.025, 95% CI [+0.006, +0.049], p = 0.008), dependence-robust, but pooled in Hz, not per SD like the rows. Signed effects positive in 17 of 28 (one-sided binomial p = 0.17); 14 of the 17 regions with |β/SE| ≥ 1 (p = 0.006). (**g**) Example sleep cycle showing the sigma-band infraslow envelope and a region-ordered spike raster. (**h**) Mean infraslow entrainment (MRL) per region. (**i**) Hourglass-encoding strength (|β_rate_|) versus infraslow-entrainment strength (MRL) for n = 1,627 units from 14 mice (log axes; Spearman ρ = 0.145, p = 4.4 × 10^−9^, pooled across animals, for display; animal-respecting tests: positive in 12 of 14 mice, p = 0.005, Wilcoxon signed-rank; animal-random-intercept mixed model β = 0.094, p = 2.3 × 10^−4^). (**j**) Per-structure coupling: raw Spearman ρ and firing-rate-controlled mixed-model β by high-level structure; the effect is significant pooled but in no single structure.

Is this macroscopic hourglass tracked by a single specialized region (a local code) or by distributed neurons (a distributed code) (**Fig. 8a**)? We tested whether single-unit firing rates during a REM episode predicted the subsequent NREM interval (|N|), using an ordinary-least-squares model adjusting for REM_pre_ duration (see Methods). β_rate_ is that unit’s ordinary-least-squires coefficient on the firing rate during REM_pre_ bouts. Only 73 of 1,609 units across the 28 regions with at least ten eligible units (4.5%, versus the 5% null expectation) passed the per-unit permutation threshold, and cohort-wide only one of 1,629 units survived FDR correction (**Extended Data Fig. 9e**). We therefore pooled units hierarchically by region. Example encoding regions are shown in **Figs. 8c,d**. Significant hourglass encoding was detected in 7 of the 32 regions eligible for the hierarchical analysis (≥ 10 units from ≥ 2 animals; **Fig. 8e**), not the 28 of the per-unit test (**Extended Data Fig. 9e**); these were dispersed across TH (4 regions), isocortex (VISp), MB (MRN) and PAL (TRS). A random-effects meta-analysis gave a positive pooled slope (restricted maximum likelihood (REML) β = 0.039, 95% CI [0.014, 0.063], p = 0.002; **Fig. 8f**). Heterogeneity was substantial (I² = 67%) and the 95% prediction interval for a new region spans zero ([−0.064, +0.141]). An animal-cluster bootstrap also gave a positive pooled slope (μ = +0.025, 95% CI [0.006, 0.049], p = 0.008, **Fig. 8f**), supporting a distributed hourglass code.

Finally, we asked what paces this distributed read-out. A natural candidate is the infraslow (0.01–0.05 Hz) oscillation of sigma-band (10–15 Hz) power, which gates NREM-to-REM sleep transitions ^15^. In our dataset, NREM-to-REM transitions occurred at a strongly preferred phase of the infraslow rhythm (**Extended Data Fig. 10**). Furthermore, this rhythm entrained single-unit population activity across the brain (**Fig. 8g,h; Extended Data Fig. 9f**).

We further examined whether the hourglass code and the infraslow rhythm reside in the same neurons, comparing each unit’s hourglass-encoding strength (|*β*_rate_|) with its infraslow entrainment (MRL) (**Fig. 8i,j**). The two were positively related: pooled across all 1,627 units from 14 mice the rank correlation accounted for ∼2% of the rank variance (Spearman ρ = 0.145, p = 4.4 × 10^−9^; **Fig. 8i**). This relationship survived the tests that respect animal identity and firing rate, holding in 12 of 14 mice individually (Wilcoxon signed-rank p = 0.005; **Extended Data Fig. 9g**). This was also the case in a mixed model with firing-rate and per-animal terms (β = 0.094, p = 2.3 × 10^−4^; **Fig. 8j**) although no single high-level structure reached significance on its own, indicating a distributed coupling rather than a localized one.

## Discussion

Here we report distributed, sleep-related neural ensembles across major structures, offering a single-cell catalogue and framing sleep as a brain-wide, coordinated process. Vigilance states reorganized firing globally with highest rates during REM sleep, mirroring previously reported vascular dynamics^61^. State was decodable from almost any region, in proportion to its neuron contribution, showing that vigilance state is globally rather than locally represented ^62^, offering potential utility for closed-loop state-dependent neuromodulation in the future ^63,64^.

Transitions were equally global, unfolding as low-dimensional trajectories spanning cortical and subcortical structures ^65,66^. At Wake–NREM boundaries, cortical populations diverged from subcortical paths ^67^, reflecting an intrinsically organized, offline cortical mode ^68^ with downstream sensory disconnection ^69^. Conversely, both REM-related transitions were unidirectional; the subcortical lead at NREM-REM transitions is consistent with parallel broadcasting from pontomedullary generators ^11,14,20,70,71^, despite diverse regional profiles ^42,72,73^. While pooling our data obscured the precise lead-lag timing between regions, these consistent cross-regional sequences ^74,75^ offer clear, feasible targets for future simultaneous recordings.

NREM slow and delta oscillations entrained spiking across nearly all regions, reflecting propagating cortical slow waves ^27,28,76^. Embedded spindles, hippocampal SWRs, and P-waves recruited distinct distributed populations ^34,77,78^ that are temporally coupled ^42,44,45^. We reported a novel temporal sequence, spindle → SWR → P-wave, extending the known nesting of slow oscillations, spindles, and ripples ^33,34^ to P-waves. How these segregated thalamocortical ^79^, hippocampal ^80^, and pontine ^39^ generators achieve this coordination remains undetermined.

REM theta rhythms entrained populations non-uniformly, consistent with localized hippocampal-limbic generators ^37,81^. However, our results reveal a brain-wide impact of P-waves, extending work tracking the brainstem-hippocampal axis ^42,44^. This distributed theta–P-wave coupling suggests that while theta entrainment is localized, phasic coordination is widespread. As P-wave-coupled theta exhibits enhanced amplitude and frequency ^40,58,82^ and theta drives memory consolidation ^83^, phasic REM sleep may distinctly modulate memory functions ^38,84,85^.

Ascending neuromodulatory anatomy partly predicted regional sleep-rhythm phenotypes. Unexpectedly, cholinergic density predicted no P-wave metrics despite its historical implication in P-wave generation ^39^, supporting recent evidence pointing to glutamatergic or medullary generators ^41^. Together with other systems, this suggests that ascending neuromodulation shapes the micro-architecture of sleep characterized by dominant oscillations during each sleep stage, although projection density is an anatomical proxy. Projection-specific manipulations may provide further insights, though optogenetic cholinergic activation suffices to induce REM sleep ^86^.

Finally, the homeostatic “hourglass” tracking REM propensity ^18^ was distributed. No single structure acted as a discrete counter; rather, a widespread signal emerged when pooled, predicting upcoming NREM accumulation and echoing the rule that REM duration forecasts the subsequent interval ^22–26,87^. This population clock co-varied with infraslow rhythms gating the NREM–REM transition ^15,35^, consistent with distributed REM-control circuits ^14,88^ and low-dimensional brain-stem gating dynamics ^16^. Whether this distributed signal participates in REM offset, or if offset is strictly governed by localized brainstem flip-flop circuits ^10,20,71^, remains to be tested.

### Limitations and outlook

Several limitations temper our conclusions. The recordings were acute and head-fixed, anatomical sampling was uneven, and REM sleep was comparatively rare. Most importantly, every relationship reported here is correlational. Resolving this will require closed-loop perturbations during defined sleep phases. Despite those caveats, a coherent picture emerges: across firing rates, transitions, oscillatory events, neuromodulatory architecture and homeostatic REM timing, the organization of sleep is distributed and coordinated. The cell-resolved atlas presented here provides the quantitative foundation on which these causal questions can be posed.

## Methods

### Animals

All animal experiments and procedures were performed in accordance with the United Kingdom Animals (Scientific Procedures) Act of 1986 Home Office regulations and approved by the Home Office (PP0688944 and PP5504676) and the University of Strathclyde’s Ethical Committee. Wild-type C57BL/6 mice of both sexes (n = 22, 10 females and 12 males, 9–28 weeks old) were used in experiments. Until surgery, the animals were housed with their littermates on a 12/12 h light/dark cycle. All recording sessions were performed at zeitgeber time 3–11. No statistical method was used to predetermine sample size.

### Surgical procedures

The procedure was done as described elsewhere ^44,89^. Mice were anesthetized with isoflurane (1–1.5%) and placed in a stereotaxic apparatus (Model 961, Kopf). To provide analgesia, Ropivacaine (Naropin, 8 mg/kg) was administered subcutaneously at the site of the incision, while Carprofen (Rimadyl, 20 mg/kg) and Buprenorphine (Vetergesic, 0.1 mg/kg) were administered subcutaneously at the back. The skin was removed from the planned head cap area, the skull periosteum was removed with 3% hydrogen peroxide, and the skull was levelled (bregma/lambda ±50 µm). Three screws (418-7123, RS Components) were implanted in the right hemisphere of the skull. Two screws were placed in the front (AP +1.5 mm, ML 1.0 mm and AP -2.0 mm, ML 2.5 mm), while the third screw was placed on the cerebellum (AP -6.0 mm, ML +2.0 mm). The posterior screw was used as a ground, while the anterior screws were used for cortical electroencephalogram (EEG) recording. Two multi-stranded wires were inserted into the neck muscle for electromyography (EMG) recording. For local-field potential (LFP) recording, a bipolar electrode was implanted in the left hemisphere pontine area (AP -5.1 mm, ML -0.6 mm, DV 3.0 mm). Finally, a custom-made semicircular headpost was attached to the skull anteriorly. During this surgery, the locations of up to 4 future craniotomy sites above the left hemisphere were also labelled. Layers of dental cement and cyanoacrylate were used to cover the skull screws and skull surface, as well as to secure the headpost in place. After surgery, mice were housed in high-topped cages with *ad libitum* access to food and water and allowed to recover for at least 5 days. After habituation to head-fixed conditions and REM sleep observation, which usually took up to 2 weeks, a second aseptic surgical procedure was done for craniotomy. As in the first procedure, mice were anesthetized with isoflurane, and analgesia was provided. Craniotomy was done over the sites marked during the previous procedure. The exposed brain surface was covered with a biocompatible sealant (Kwik-Sil, World Precision Instruments). The first Neuropixels recording session followed the next day.

### Electrophysiological recording

Neural population activity was recorded in 4 hr daily sessions over up to 4 days. Head-fixed, awake mice were set in a restraining tube within a soundproof box (MAC-3, IAC Acoustics) in dim light conditions. At the start of each recording session, Kwik-Sil was removed, and 1% agar was used to protect the brain surface. Electrophysiological recordings were done with Neuropixels 1.0 and 2.0, mounted on a manipulator (DMA-1511, Narishige or Neuropixels mini-compact unit, Luigs&Neumann). Probes were inserted stereotaxically, in the pre-planned location and trajectory angle, based on mouse brain atlas measurements. Insertion speed was 10–100 µm/s, applied intermittently, and signals were checked throughout the implantation. Probes used external reference and ground, connected through a soldered wire to a mouse head cap ground point. Signals collected through the Neuropixels probe were amplified and digitized in the probes integrated circuit and recorded at a 30 kHz sampling rate using SpikeGLX (Janelia Research Campus). The recording sessions were typically initiated approximately 1 hr after the probe insertion, to allow for signal stabilization. The rear of the probe was painted with CM-DiI (C7001, Invitrogen, 0.1% w/v diluted in ethanol) before insertion, allowing us to identify the probe tracks histologically. Cortical EEG, EMG and pontine LFPs were amplified (HST/16 and PBX3, Plexon), digitized and recorded at a 2.5 kHz sampling rate using SpikeGLX.

### Face video recording and processing

A camera (acA1920-25um, Basler Ace) with a zoom lens (M0814-MP2, Computar) and an infrared filter (FGL780, Thorlabs) was used to monitor and record facial movements of animals during recording sessions. Images were collected at 25 Hz using a custom-written LabVIEW program and a National Instruments image grabber (PCI-6221). A new video file was saved every 60 sec, and for each new file a start sync pulse was generated and sent to SpikeGLX for electrophysiology-to-video data alignment.

Video files were processed using DeepLabCut (DLC 2.3) ^90^. For pupil tracking, four keypoints were labeled: left, right, top and bottom edge of the pupil. Training dataset consisted of 400 frames extracted across 10 videos from different mice. Further, a confidence threshold was applied to DLC filtered output data, where only the frames where all four key-points exceeded a likelihood of 0.6 were retained, and the rest were excluded. To compute pupil diameter, the lateral diameter was calculated across frames. To compute eye movement, the Euclidean distance of the centre point (midpoint between the left and right edges) was calculated between consecutive frames.

### Histology

After completing recording sessions, mice were injected intraperitoneally with pentobarbital (200 mg/ml) and transcardially perfused with 0.1M PBS and 4% paraformaldehyde in PBS. Brain tissue was extracted and stored in the same fixative overnight at 4°C, then transferred into 30% sucrose solution for at least 2 days. Tissue was cut into 80 µm coronal sections using a microtome (SM2010R, Leica), stained with DAPI (1:1000, #62248, ThermoFisher Scientific), and imaged for CM-DiI probe tracks under an epifluorescent upright microscope (Eclipse E600, Nikon) or an automated slide imaging system (EVOS M7000).

### Probe track registration

Probe channels registration was performed as described previously ^49,89^. SHARP-Track (https://github.com/cortex-lab/allenCCF/tree/master/SHARP-Track) was used to register histological slices in the Allen 3D atlas, and label the probe track location (**Extended Data Fig. 2a**). SHARP-Track output gave the list of brain regions and their estimated borders in µm. For Neuropixels 1.0 probes, where the whole length of the shank bank was sampled, the Kilosort-processed multi-unit activity was plotted and assessed across probe channels, and specific brain region borders were identified based on their spiking patterns (**Extended Data Fig. 2b**): for example, the cortical surface and white matter are easily identifiable through a sudden drop in spikes on both sides. The locations of these identified borders in Neuropixels channels were then cross-compared with the histologically estimated border locations (**Extended Data Fig. 2c**). Median error between electrophysiologically and histologically estimated borders across datasets was 78 µm (**Extended Data Fig. 2d**), which is comparable with previous reports ^49,89^. Combining SHARP-Track information on sampled brain areas with known brain-area specific MUA activity allowed us to determine which recording channels were located in which brain regions.

For Neuropixels 2.0 probes, where channels selected for recording were spread across the tips of 4 shanks, we could not use brain region borders to assess their locations because all selected channels were located within a few brain regions without clear electrophysiological markers. Instead, we relied on stereotaxically measured insertion depth, and identification of the CM-DiI probe tips in histological brain slices. SHARP-Track was still used in this case for identification of brain regions, and their location in µm was matched to measured insertion depth.

### Preprocessing of Neuropixels signals – single units

Preprocessing to isolate single units consisted of the following four steps (**Extended Data Fig. 3a**).

1. **Spike sorting.** Raw extracellular voltage traces from Neuropixels probes were spike-sorted with Kilosort4 ^91^. The pipeline applied a 300 Hz high-pass filter, common-average referencing across channels, and non-rigid drift correction estimated from continuous spike-feature tracking. Spikes are then detected and assigned to clusters by iterative template matching, with templates re-estimated on each pass and automated splitting and merging applied at the end. Sorting was run with the default Kilosort4 parameters, producing a per-recording set of putative clusters together with their template waveforms, peak channel, mean depth, and spike times. All subsequent quality control was performed post-hoc on these Kilosort4 outputs. Given the volume of the data and for reproducibility, no manual curation was applied beyond the automated steps described below.
2. **Region assignment.** Each cluster was assigned to its peak Kilosort4 channel and the channel position was mapped to the Allen Mouse CCF v3 ^53^ using per-recording probe-trajectory measurements described above. The CCF region closest to the channel coordinate was recorded as the cluster’s final region. Clusters whose nearest annotation fell on fiber tracts, ventricles, the root node, or any other structure not belonging to one of the nine high-level structures were flagged as anatomically ineligible and excluded from all subsequent analyses. These clusters included border regions that CCF v3 did not assign any specific region that typically appears in the midbrain and the pons in our datasets.
3. **Cluster quality control.** Each Kilosort4 cluster was independently classified using Bombcell v0.70 (Python) (https://github.com/Julie-Fabre/bombcell) ^92^, an open-source Python toolbox that computes a panel of waveform-shape and firing-statistics metrics and applies a deterministic decision tree to assign every cluster to one of four mutually exclusive categories: NOISE, multi-unit activity (MUA), well-isolated single unit (GOOD), or non-somatic (NON-SOMA). Only GOOD clusters were included.
4. **Region-level inclusion filter**. To avoid drawing inferences from sparsely sampled brain regions, a region-level inclusion filter was also applied on top of the per-cluster criteria described above (**Extended Data Figs. 3b-d**). A brain region was retained only if (i) it contributed at least 10 GOOD units pooled across recordings and (ii) was sampled in at least two recordings. Depending on analysis types, additional filters were applied.

### Preprocessing of Neuropixels signals – LFP

LFP signals were extracted from each Neuropixels probe by downsampling lowpass signals from Neuropixels 1.0 probes or broadband signals from Neuropixels 2.0 probes at 1 kHz. Half of the active channels (192) were retained so that the full span of the recorded area was preserved. For every recording, each retained channel was projected back onto its physical position on the probe and assigned a brain region by nearestneighbor matching to the spike-sorted clusters whose anatomical regions were determined in the Allen Mouse CCF.

### Sleep scoring

States were visually scored offline, as described elsewhere ^42,44^. Wakefulness, NREM, or REM sleep was determined for every 4-second window based on cortical EEG and EMG signals using a custom-made MATLAB GUI (https://github.com/Sakata-Lab/SleepScore). Wake was characterized by high EMG power and low EEG delta power. NREM sleep was characterized by high EEG slow/delta power, low EEG theta power, and low EMG power. REM sleep was characterized by high EEG theta power, low EEG delta power, and low EMG power. The same individual scored all recordings for consistency.

### Firing rate comparison

Spikes were assigned to non-overlapping 4-second epochs whose vigilance-state label (NREM, REM or Wake) was scored as described above. State-specific mean firing rates were calculated by dividing total spike counts across all epochs of a given state by that state’s cumulative duration. Neurons with fewer than five epochs (i.e., 20 s) in any of the three states were excluded from subsequent analyses.

Neuron-wise state modulation was assessed using a two-stage permutation approach on epoch-level firing rates. First, an omnibus one-way F-statistic computed across states was compared against a null distribution generated from 10,000 random epoch-label permutations. Neurons failing this test (*p* ≥ 0.05) were classified as Neutral. For the remaining state-modulated cells, three pairwise permutation tests evaluated differences in mean firing rates, yielding p-values and signed log_2_ fold-changes. Units were then categorized into seven mutually exclusive classes based on these pairwise outcomes: (i) Single-state preferred (REM-, NREM-, or Wake-preferred): Both pairwise comparisons involving that state were significant (*p* < 0.05) and consistent in sign. (ii) Bistate (REM/NREM-, REM/Wake-, or NREM/Wake-bistate): Firing rates between two states were statistically indistinguishable, but both differed significantly (and in the same direction) from the third state. (iii) Neutral: Failed the omnibus test or did not match single-state or bistate criteria. To control the cohortwide false discovery rate (FDR) across thousands of units, the Benjamini-Hochberg (BH) procedure was applied at *q* = 0.05 independently to each of the four p-value families (one omnibus, three pairwise) pooled across all units.

To examine regional differences, we calculated the mean and standard error of per-state firing rates across clusters for each high-level anatomical structure. Two complementary statistical tests were applied: (i) Within-structure comparisons: Pair-matched within-cluster state differences were evaluated using Wilcoxon signed-rank tests on cluster-wise log_10_-transformed firing rates (with a 0.01 Hz floor added prior to transformation). The resulting p-values were BH-corrected across all structures. (ii) Across-structure comparisons: To capture the global direction of the contrast at the anatomical level, an additional Wilcoxon signed-rank test was performed across the nine high-level structures. Region-level contrasts are one-sample Wilcoxon signed-rank tests on the per-region log-fold differences (n = 43 regions).

### Vigilance state decoding

A recording was used only if it met the following three criteria: (i) at least ten quality-passing neurons, (ii) at least thirty epochs (120 s) of each of the three vigilance states, and (iii) at least one REM episode of any length. For each retained recording, a population firing-rate matrix was constructed at the 4-second resolution. Each bin was the firing rate (Hz) of one neuron in one epoch. Firing-rate features were z-scored prior to classifier training, and standardization was performed strictly within each cross-validation fold: the mean and standard deviation used to transform a held-out test partition were estimated exclusively from the corresponding training partition.

#### All-neuron random forest (RF) decoder

Vigilance state was decoded from the population firing-rate matrix with a Random Forest (RF) classifier configured with 500 decision trees, no maximum depth constraint, a minimum of five samples per leaf node, and class weights set inversely proportional to class frequency. Each 4-second epoch was treated as one labelled sample whose feature vector was the firing rates of all neurons recorded in that recording during that epoch. Because class imbalance was intrinsic (e.g., REM was comparatively scarce), the inversely frequency-weighted class weights up-weighted minority classes during tree construction so that no class was systematically favored.

Decoder performance was evaluated by 5-fold temporal-block cross-validation. The primary performance metric was balanced accuracy, defined as the unweighted mean of per-class recall values. The contribution of each neuron to the all-neuron decoder was quantified by the Gini-based mean decrease in impurity (MDI). For a tree node *t* holding training samples of the three vigilance-state classes (*K* = 3), the Gini impurity is

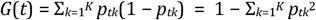

where *p_tk_* is the class-weighted proportion of samples of class *k* reaching node *t*; because the forest used class weights set inversely proportional to class frequency, these proportions are weighted so that the scarce REM class is not discounted. When node *t* is split into a left and a right child (*t*_L_, *t*_R_), the impurity decrease attributed to that split, weighted by the fraction of samples reaching the node, is

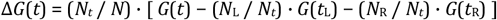

with *N_t_*, *N*_L_ and *N*_R_ the (weighted) numbers of samples reaching node *t* and its left and right children, and *N* the total at the root. The raw importance of feature *f* in a single tree *j* is the sum of these decreases over every internal node whose split variable *v*(*t*) selects *f*,

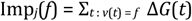

Each tree’s importances are normalized to sum to one and then averaged over the *T* = 500 trees of the forest, giving the per-neuron importance reported for one cross-validation fold,

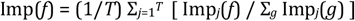

so that the importances are non-negative and sum to one across all neurons of a recording (note that impurity-based Gini importance is biased towards correlated and high-cardinality predictors; unless stated otherwise, relative Gini importance, defined below, is the measure reported in the main figures). This per-fold importance was averaged over the five cross-validation folds to yield a single value per neuron,

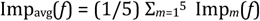

To make importances comparable between recordings of differing neuron yield, each averaged value was multiplied by the number of neurons entering the decoder in its own recording,

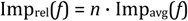

where *n* is that recording’s neuron count, so that a value of one denotes a neuron of exactly average importance within the model that was actually fitted.

Region-level importance was then obtained by averaging the relative importances of the neurons assigned to each anatomical region *r*,

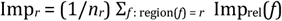

where *n_r_* is the number of neurons assigned to region *r*.

#### Region-specific RF decoders

To quantify how much vigilance-state information was carried by each anatomical area independently of the rest of the recorded population, a separate RF classifier was trained for every brain region using only neurons located within that region. Classifier hyperparameters, the cross-validation scheme, the standardization procedure and the performance metrics matched those of the all-neuron decoder exactly; only the feature space changed. Regions contributing fewer than three neurons in a given recording were excluded from this analysis.

#### Permutation testing

Statistical significance of every decoder was assessed by a non-parametric permutation test. Vigilance-state labels were shuffled with independent random number generators, and the full cross-validation procedure was repeated on each shuffle to construct an empirical null distribution of balanced-accuracy values. The all-neuron decoder used 500 permutations; region-specific decoders used 200, a reduction adopted to keep the wall time tractable given the substantially larger number of region-by-recording models. Null forests were matched to the observed decoder in all hyperparameters, including tree count. The permutation p-value was computed with the add-one estimator, p = (1 + Σ)/(1 + B), where Σ is the number of null balanced accuracies at or above the observed value and B the number of shuffles (one-sided), so the attainable floor is 1/501 for the all-neuron decoder and 1/201 for the region-specific decoders. Region-restricted p-values were Benjamini–Hochberg-corrected across the 181 region × recording tests, and a region was additionally required, under the stricter all-recordings criterion, to be significant in every recording sampling it.

### State transition analysis

#### Matrix construction and static principal component analysis (PCA)

For each brain region *i*, a peri-transition firing-rate profile *r_i_*(*t*) was built as follows. Spikes from all eligible neurons in that region (mean firing rate ≥ 0.1 Hz) were pooled and binned at 10 ms over a ±60 s window around every scored transition. Each bin was then restricted to the canonical pre- or post-transition state, so that only epochs whose scored state matched the expected one contributed (pure-state masking). The profile was smoothed with a NaN-aware Gaussian convolution (FWHM = 2 s) and z-scored against the [−60, −30] s baseline window. Static PCA on this matrix yielded a single PC1–PC2 point per region.

#### Delay-embedded PCA

The PCA described above discards the within-window temporal evolution of the firing-rate profile. To recover this dynamic information, a delay-embedded variant of the analysis was performed. For each region, the profile was unfolded into a Hankel-like matrix *H_i_* of size (*C* × *L*) whose *k*-th row contained the contiguous lag-vector

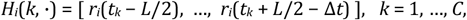

with window length *L* = 4 s = 400 bins and *C* = 121 window centers *t_k_* spaced by *h* = 0.5 s and tiling the display interval [−20, +40] s. Stacking across the *R* sampled regions gave the joint embedding matrix

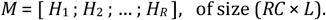

Each of the *L* lag features (columns of *M*) was standardized to zero mean and unit variance (scikit-learn *StandardScaler*), and the resulting matrix decomposed by singular value decomposition (scikit-learn *PCA*),

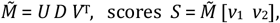

with the two leading right-singular vectors taken as PC1 and PC2. The fraction of variance explained by component *k* was *λ_k_* / Σ*_j_ λ_j_*, with *λ_k_* = *σ_k_*^2^ / (*RC* − 1), where *σ_k_* is the *k*-th singular value (the *k*-th diagonal entry of *D*).

To quantify the qualitative cortical-versus-subcortical separation, every region was assigned to one of two anatomical groups on the basis of its CCF high-level structure: a “cortical” group comprising the isocortex, hippocampus and lateral amygdala (LA), and a “subcortical” group comprising the remaining structures. For each region, a displacement vector running from a pre-transition centroid (the mean of its trajectory between −20 and −5 s) to a post-transition centroid (the mean of its trajectory between +5 and +20 s) was computed. Each displacement vector was characterized by its angle in the PC1–PC2 plane and by its Euclidean magnitude. The angular distributions of cortical and subcortical regions were compared with the Watson–Williams test (a circular analogue of one-way analysis of variance), with the angular difference between the two group means and the test p-value annotated on the polar histogram of each transition. The magnitude distributions were compared between cortical and subcortical regions with the Mann–Whitney U test, with the test p-value annotated on the boxplot of each transition. Recording-level bootstrap resampling (1000 iterations) was used throughout to obtain 95% confidence intervals on group means.

### Sleep-related event detection

#### Sleep spindles

Sleep spindle detection was done based on a previously published approach, with parameters optimized for our recordings ^93^. Cortical EEG was bandpass filtered at 8–16 Hz using a 4th-order Butterworth filter for subsequent cycle counting. Independently, a continuous Morlet wavelet transform was applied to the raw EEG, restricted to the same frequency range. The squared magnitude of the wavelet coefficients was averaged across frequencies to generate a single spindle energy envelope, which was smoothed using a 200 ms Hann window. Detection thresholds were calculated from the smoothed energy envelope in NREM epochs, and candidate events had to pass the high threshold (mean + 2.5 SD) for event detection. This was followed by the low threshold (mean + 0.8 SD) to find the start and end of detected events. There were additional validation criteria for an event to be considered a sleep spindle: (1) Event duration between 0.4 and 2 seconds, (2) Event containing between 5 and 30 cycles, estimated as half the number of zero crossings in the bandpass-filtered signal, (3) Power in the spindle band (8–16 Hz), calculated from the raw EEG over the candidate event, had to exceed power in both adjacent bands (6–8.5 Hz and 16.5–20 Hz), and (4) No overlap with a previously accepted spindle.

#### Sharp-wave ripples

Sharp-wave ripples (SWRs) were detected from the 1 kHz sub-sampled LFP signal of each Neuropixels probe. Recordings were retained when the probe sampled at least one channel assigned to the hippocampal CA1 field and the four-second sleep score contained both NREM and REM epochs. Within every eligible recording, the SWR detection channel was selected as the within-CA1 maximum of the NREM-to-REM ripple-band (140–250 Hz) power ratio, exploiting the strong suppression of ripple-band activity at the pyramidal layer during REM. The LFP trace of the selected channel was then bandpass filtered between 140 and 250 Hz with a third-order Butterworth filter applied bidirectionally, rectified and smoothed with a 20 ms root-mean-square sliding window to give a ripple-band envelope. NREM-conditioned envelope thresholds of mean plus five standard deviations (peak threshold) and mean plus two standard deviations (boundary threshold) defined candidate events; overlapping candidates were merged and events whose duration fell outside 20 to 200 ms were discarded. To suppress movement-artefact contamination of the ripple pool, every candidate event was then validated against the EMG recorded simultaneously with the session. The EMG root-mean-square across each event’s onset-to-offset window was compared with the 95th percentile of the per-recording distribution of NREM-event EMG root-mean-square values, and events exceeding that threshold were tagged as artefactual and excluded from all downstream analyses.

#### Pontine waves

P-waves were detected from two complementary signal sources in this study: bipolar EEG electrodes implanted into the pons, and bipolar derivations constructed offline from neighboring channels of a Neuropixels probe. The two pipelines share the same underlying logic ^42,44^: (1) subtract two nearby pontine signals to remove common-mode activity, (2) band-pass filter, and (3) detect negative deflections that exceed a noise-derived threshold.

For P-wave detection from pontine bipolar electrodes, the two channel signals were subtracted to obtain the differential pontine signal, which was band-pass filtered between 5 and 30 Hz. To set the detection threshold, a noise segment was taken from the longest NREM sleep episode of the recording, and the root-mean-square (RMS) of the filtered signal was computed in non-overlapping 10 ms windows; the threshold was defined as the mean of these RMS values plus five standard deviations. Threshold crossings were treated as P-wave candidates, and the timing of each P-wave was assigned to the negative peak of the local deflection. To suppress movement artifacts, EMG RMS was computed with the same 10 ms resolution and any candidate whose EMG RMS exceeded the EMG mean plus three standard deviations was discarded.

For P-wave detection from Neuropixels LFPs, eligible channels were first restricted to the mesopontine regions LDT, PPN and PB, which an initial channel-optimization analysis on the cohort identified as the regions in which canonical P-waves were most clearly resolvable (**Extended Data Fig. 7**). From this candidate pool, channel pairs were enumerated and ranked first on whether both channels fell within the preferred-region set, and second on a signal-to-noise proxy equal to the standard deviation of the 10 ms RMS values divided by their mean, computed on the longest NREM episode. The optimal inter-electrode distance was constrained to 500 ± 50 µm, with a documented fallback to the wider 200–800 µm range when no pair satisfied the narrower window (**Extended Data Fig. 7**). The trace was band-pass filtered between 5 and 30 Hz with a 5th-order Butterworth filter, applied bidirectionally so that no phase delay was introduced. A noise calibration segment was extracted from the longest NREM episode (clipped to between 8 and 48 s when necessary), and the RMS of the filtered signal was computed in non-overlapping 10 ms windows. The detection threshold was the mean of these RMS values plus five standard deviations, negated. It is therefore equal in magnitude to the threshold used for the pontine bipolar electrodes above, but signed so that it can be applied directly to the raw filtered trace, which carries the negative-going P-wave deflection. Each threshold crossing initiated a search for the local negative peak, taken as the argmin over the 50 ms window starting at the crossing. The sample of that minimum was registered as the P-wave time, and the surrounding waveform was stored for subsequent quality control. A 50 ms refractory period was then enforced before the next search resumed.

Two artifact-rejection passes were then applied. First, EMG RMS was computed across the entire recording in the same 10 ms non-overlapping windows, and candidate events whose enclosing EMG-RMS window exceeded the EMG mean plus three standard deviations were discarded as putative movement artifacts. Second, the peak-to-trough amplitude was measured on each stored candidate waveform; a reference distribution of peak-to-trough amplitudes was built from the REM-state subset of candidates (or, when too few REM events were available, from NREM candidates), and any event whose peak-to-trough exceeded the reference mean plus three standard deviations was removed as an outlier.

### Oscillation phase analyses

#### Spike-phase analyses

The frontal cortical EEG was independently bandpass filtered into a 0.02–0.5 Hz slow band, a 0.5–4 Hz delta band and a 6–10 Hz theta band using a Butterworth filter applied bidirectionally so that no phase delay was introduced. The instantaneous phase of each filtered band was then obtained as the angle of the analytic signal produced by the Hilbert transform. This yields a continuous phase trajectory in which zero phase corresponds to the positive peak of the band-passed waveform and ±π to the trough.

For every eligible neuron, each spike was assigned the instantaneous EEG phase of its enclosing time bin, interpolated linearly from the milli-second-resolution phase trace, separately for each band: spikes during NREM sleep for the slow and delta bands, and spikes during REM sleep for the theta band. The mean resultant length (MRL) of the spike-aligned phase distribution was computed as the modulus of the complex average of the unit vectors exp(i*φ_n_*) over the *N* spikes of the unit. The preferred phase of the unit was the circular mean of the same distribution, and a Rayleigh test of non-uniformity was applied to call each unit significantly phase-locked or not at a significance level of 0.05.

#### NREM sleep event phase analyses

Event phases were obtained in the same way. The frontal cortical EEG was bandpass filtered into the same slow (0.02–0.5 Hz) and delta (0.5–4 Hz) bands and its instantaneous phase extracted by the Hilbert transform. For every event, the EEG phase at the event reference time was interpolated linearly from the millisecond-resolution phase trace, giving a per-session vector of event-aligned phases for each event type and band. The MRL of that vector was taken as the depth-of-modulation metric. Group differences across the three event types within a band were assessed by a Kruskal-Wallis omnibus test on the across-session distributions of the MRL.

### NREM sleep event coupling

The pairwise temporal coupling among the three NREM events was quantified with two complementary descriptors. The first was the cross-correlogram. For every pair of event types and every session, the lag from each reference event to every target event was binned into 25 ms bins over a ±1 s window and normalized by the number of reference events, giving a rate-per-event histogram. The second descriptor was the nesting rate, in which for each event pair and each fixed nesting window of ±100, ±250 or ±500 ms we counted the number of target events that fell within the window of every reference event and divided by the number of reference events. A circular-shift shuffle null distribution was used throughout. Shuffling event times across the whole recording would spuriously disperse them onto REM and Wake intervals and bias every null distribution downward, so we first concatenated the NREM segments of each session into a single continuous stitched NREM-only timeline. Every event time was remapped onto this stitched axis, and the target event times were then circularly shifted on the stitched axis by a random offset on each of one thousand shuffles for the cross-correlograms and five hundred for the nesting rates. Each shuffled event is therefore guaranteed to remain in NREM by construction, so the resulting null distribution reflects the geometry of the NREM timeline rather than the alternation of vigilance states. The shuffle distribution gives an empirical 95% confidence band that is overlaid on every observed cross-correlogram, and a shuffle p-value for every observed nesting rate.

### Theta-P-wave-coupled spike analysis

For every eligible unit that fired at least 100 spikes during REM, every P-wave was first labelled with the cortical theta phase at the P-wave’s negative peak and binned into 8 theta-phase bins spanning the cycle from trough through peak to trough. A peri-event firing-rate histogram was then computed separately for each phase bin. The unit’s spike times were aligned to the P-waves whose theta phase fell into that bin and binned at 25 ms over a ±2 s window. The histogram was smoothed with a Gaussian kernel of two-bin standard deviation and z-scored against the −2.0 to −1.5 s pre-event baseline used elsewhere. The eight phase-stratified histograms together form a phase × time matrix per unit, which was averaged across the units of each Allen CCF region to give a region-level phase × time map; regions with fewer than 5 contributing units were dropped from the cohort test.

Phase-gating of the P-wave response was then quantified per region by a non-parametric Friedman test on the per-unit mean post-event z-score in the response window (±0.1 s around the P-wave) across the 8 phase bins, treating each unit as a repeated measure. The Friedman chi-squared statistic was converted to a normalized effect size using Kendall’s coefficient of concordance:

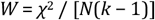

where *N* is the contributing unit count and *k* = 8 is the number of theta-phase bins, so that effect sizes are comparable across regions of different unit counts. Per-region Friedman p-values were Benjamini-Hochberg corrected across all regions.

### Neuromodulator projection maps

Whole-brain axonal projection strengths for each mesopontine neuro-modulator system were obtained from the Allen Mouse Brain Connectivity Atlas through the AllenSDK Mouse Connectivity Cache. Datasets for four neuromodulator systems were extracted: dopaminergic (*Slc6a3*-Cre), serotonergic (*Slc6a4*-CreERT2), noradrenergic (*Dbh*-Cre KH212), and cholinergic (*Chat*-IRES-Cre). Cre-gated systems were restricted to experiments whose injection envelope covered the canonical source nucleus of that cell type: VTA/SNc/SNr for dopamine, dorsal raphe for serotonin, locus coeruleus for noradrenaline, and the pedunculopontine and laterodorsal tegmental nuclei (PPN/LDT) for acetylcholine (n = 2, 2, 3 and 2 experiments, respectively). Primary Allen injection-structure labels coincided with that nucleus except for noradrenaline, whose labels are injection centroids in the laterodorsal tegmental or parabrachial nucleus rather than locus coeruleus (**Extended Data Fig. 8**); the noradrenergic map should therefore be read as reflecting LC-territory rather than LC-exclusive innervation. For each qualifying tracer experiment, we extracted the normalized projection volume - the projection signal summed within an Allen CCF target region and divided by the source-injection volume and averaged this value across experiments to give a single projection strength per neuromodulator system and target region. The mean was log-transformed prior to analysis.

### Projection–phenotype correlation

For each combination of neuromodulatory innervation and electrophysiological metric, we assessed the across-region association between log mesopontine projection strength and the metric using the two-sided signed Spearman rank correlation (ρ), computed over the CCF regions for which both quantities were defined. A minimum of three regions was required for a given system-by-metric combination to enter the analysis. Significance was reported with family-wise false-discovery-rate correction.

### Joint embedding and feature smoothness

To examine how regions are organized jointly by their innervation and their physiology, a single feature matrix was constructed with ten standardized variables: the four log mesopontine projection strengths and the six state-dependent neurophysiological metrics. Regions missing any feature were excluded, and each feature was z-scored so that all entered the analysis on a common scale. The resulting region-by-feature matrix was embedded in two dimensions using Uniform Manifold Approximation and Projection (UMAP) with a Euclidean metric, a neighborhood size of ten, a minimum distance of 0.30 and a fixed random seed for reproducibility.

Because UMAP is non-linear and its axes carry no interpretable loadings, we quantified the contribution of each feature to the embedding by how smoothly that feature varies over the neighbor graph that UMAP itself constructs. For each feature we computed Moran’s I on UMAP’s fuzzy weighted graph,

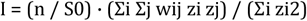

where n is the number of regions, wij is the weight of the graph edge between regions i and j, S0 is the sum of all edge weights, and zi is the mean-centered standardized value of the feature at region i. The graph scored here is the fuzzy neighbor graph that UMAP constructs in the ten-dimensional feature space; the two-dimensional embedding itself never enters the statistic.

Because that graph is built from the same ten features that are then scored on it, Moran’s I is positively biased and its null expectation lies well above zero. Every feature was therefore calibrated against a permutation null that preserves this dependence: the feature’s values were permuted across regions, the fuzzy neighbor graph was rebuilt from the nine intact features together with the permuted one, and Moran’s I of the permuted feature was recomputed on that rebuilt graph (10,000 permutations per feature). Null means obtained this way were feature-specific, ranging from 0.196 to 0.342, so no single global baseline is appropriate. Each observed value was referred to its own null distribution to give a two-sided empirical p value (add-one corrected), and these were converted to Benjamini–Hochberg q values across the ten features. Because this null is itself conditional on the other nine features, we additionally scored each feature on a graph built from the other nine alone; the weight matrix is then independent of the feature under test, so the classical permutation null applies exactly. A feature is reported as exceeding chance only where it clears both designs at q < 0.05, which five of the ten features do (**Fig. 7j**). Two further features clear the second design only; their status depends on which null is specified, and no claim is made about them in either direction.

### Hourglass model and infraslow oscillations

#### Behavioral hourglass regression

A REM cycle was defined as a triplet consisting of an opening REM bout, REM_pre_, an inter-REM interval, and a closing REM bout, REM_post_. |*N*| was computed as the total time spent in NREM sleep within the inter-REM interval. To prevent arousal-fragmented sleep from being treated as a single hourglass cycle, any cycle whose longest contained Wake bout exceeded 120 s was excluded. The pooled hourglass regression was fitted by maximum likelihood as a linear mixed model with a per-animal random intercept and a random slope for REM-bout duration, to absorb between-subject differences in both baseline and sensitivity: log|*N*| ∼ log(REM_pre_) + (1 + log(REM_pre_) | animal).

#### Per-cycle single-unit firing-rate regression

For each unit, the firing rate during REM_pre_ period was computed for each of that unit’s eligible cycles. |*N*| was then regressed on this firing rate by ordinary least squares, adjusting for the log REM-bout duration in the same model (log|*N*| ∼ log(REM_pre_) + rate), so that the per-unit rate coefficient *β*_rate_ reflects firing variation beyond what REM-bout duration alone predicts. Significance of the rate coefficient was assessed against a within-recording permutation null. The pairing between cycle firing rates and inter-REM durations was randomly shuffled 500 times within the same recording, preserving recording- and animal-level structure, and the model was refitted on each shuffle. The two-sided empirical p-value is the fraction of shuffles whose rate coefficient was at least as large in absolute value as the observed one. Because most units contributed only a handful of cycles, this permutation null has a coarse, high effective floor: false-discovery-rate correction is unattainable for the great majority of units even when an underlying effect is real, and regional yields of per-mutation-significant units therefore cluster near the 5% null expectation. This was why the hierarchical pooling was taken.

#### Hierarchical per-region mixed model

To recover the regional signal that the single-unit test cannot resolve, we pooled each unit’s cycles within its Allen CCF region. Before pooling, each unit’s per-cycle firing rate was standardized to zero mean and unit variance within that unit (rate_z_), so that stable between-unit differences in baseline rate could not contaminate the region-level slope; rate_z_ therefore carries only each unit’s cycle-to-cycle firing fluctuations. A region was analyzed only if it contributed at least ten eligible units from at least two animals. Within each qualifying region we fitted a linear mixed model

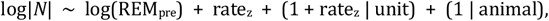

in which the log |*N*| was predicted jointly by the log REM_pre_ duration and the standardized firing rate, with a random intercept and a random rate_z_ slope per unit and a random intercept per animal (entered as a variance component). The fixed-effect slope on rate_z_, denoted *β*, was the region-level coupling between single-unit REM firing and |*N*|. Significance was assessed against a within-recording permutation null in which rate_z_ was shuffled within recording (200 shuffles per region, refitting the mixed model each time; two-sided empirical *p* as above), and the resulting per-region *p*-values were corrected across all regions tested with the Benja-mini–Hochberg procedure at *q* = 0.05.

#### Distributed-signal random-effects meta-analysis

Because counting individually FDR-significant regions cannot distinguish a focal signal from a distributed one that is broadly present but individually under-powered, we treated each region as a single observation, parameterized by its slope *β*_i_ and standard error *SE*_i_, and pooled the regions by random-effects meta-analysis. Inverse-variance weighting requires a finite *SE*_i_, and four of the 32 regions (LA, LSc, PPN and TRS) returned a singular random-effect covariance and so had none, leaving *k* = 28; all 32 are plotted in **Fig. 8f**, the four excluded ones without a confidence interval. The pooled mean is the inverse-variance-weighted average

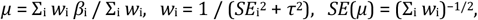

with a Wald 95% confidence interval and the between-region variance *τ*^2^ estimated by iterative restricted maximum likelihood (REML) ^94^. Heterogeneity is reported as Cochran’s *Q*, *I*^2^ and a 95% prediction interval. Because regions that share an animal are not independent, the magnitude of the pooled effect was assessed by an animal-cluster bootstrap over the 14 animals of the unit-level slope table (2,000 iterations, resampling whole animals with replacement and refitting the REML pool on each) and its breadth by a one-sided exact binomial sign test on the signed regional slopes, reported both over all *k* regions and over the subset with |*β*_i_ / *SE*_i_| ≥ 1.

#### Infraslow sigma-power phase estimation

To relate firing to the NREM sigma-power infraslow oscillation, a polarity-robust infraslow phase from the cortical EEG was derived, following the previously described sigma-envelope approach ^35,36^. The EEG was band-pass filtered to the sigma band (10–15 Hz) with a fourth-order zero-phase Butterworth filter (second-order sections, forward–backward filtering), and the sigma-band amplitude envelope was taken as the modulus of the Hilbert analytic signal. The envelope was decimated to 10 Hz, band-pass filtered to the infraslow band (0.01–0.05 Hz) with a second fourth-order zero-phase Butterworth filter, and Hilbert-transformed to yield a continuous infraslow phase φ(*t*) in (−π, π], with 0 radians at the sigma-envelope peak. Because the envelope is unsigned, this phase is invariant to EEG polarity. All entrainment analyses were restricted to NREM: a time point was treated as NREM if the 4-s epoch containing it was scored NREM.

#### Single-unit phase locking and regional entrainment

For each unit, every NREM spike was assigned the infraslow phase at its time of occurrence by interpolating φ(*t*) (sampled at 10 Hz) at the spike time. The strength of phase locking was summarized by the MRL,

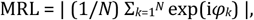

where the sum runs over the *N* infraslow phases *φ*_k_ of that unit’s NREM spikes (MRL ranges from 0 for a uniform phase distribution to 1 for perfect phase locking). Units with fewer than 100 NREM spikes were not evaluated. Statistical entrainment was judged against a circular time-shift surrogate null that preserves each spike train’s autocorrelation while destroying its phase alignment: the spike train was rigidly shifted by a random offset drawn uniformly from [60 s, *T* − 60 s] (where *T* is the recording duration) and wrapped modulo *T*. The NREM mask was then re-applied, the infraslow phase re-interpolated and the MRL recomputed. This was repeated for 200 surrogates. An entrainment *z*-score, *z*_MRL_, was formed as

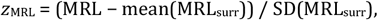

and a unit was classified as entrained when *z*_MRL_ ≥ 2. The parametric Rayleigh-test false-discovery-rate flag was degenerate at this spike count (it passed essentially all units) and was not used for selection.

#### Coupling between hourglass encoding and infraslow entrainment

To evaluate the relationship between hourglass-encoding strength and infraslow entrainment, each unit’s absolute per-unit rate coefficient (|*β*_rate_|, the ordinary-least-squares slope defined under “Per-cycle single-unit firing-rate regression”, in units of log|*N*| per Hz) was matched to its MRL by joining the two datasets on recording folder and cluster identity; 1,627 units from 14 mice carried both measures. Because both encoding strength and entrainment covary with baseline firing rate, a linear mixed-effects model (LMM) was used to control for confounding firing-rate effects and to account for inter-animal variability. |*β*_rate_| and the unit’s mean firing rate over the recording (FR) were log_10_-transformed (values of |*β*_rate_| below 10^−4^, of which there was one, were floored at 10^−4^) and then, with the MRL, standardized to zero mean and unit variance across the units entering each fit; *z*(·) denotes that standardization. The primary model was defined as:

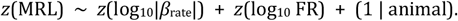

where (1 | animal) represents a random intercept per animal. To test the robustness of the fixed effect, a secondary model adding a per-animal random slope for *z*(log_10_|*β*_rate_|) was also evaluated. Finally, to assess whether the relationship was anatomically localized or a brain-wide property, this primary LMM framework was fitted independently within each Allen high-level structure that contributed at least 30 units from at least three mice, with all variables re-standardized within that structure; structures below that threshold are reported with the firing-rate-partialled Spearman correlation only.

### Statistics and reproducibility

#### Unit of statistical inference

Ten units were used across this hierarchical dataset: neuron, Allen CCF region, high-level CCF structure, recording, animal, sleep cycle, detected event (spindle, ripple or P-wave), electrode pair, 4-s epoch and region × recording decoder. The unit for each of the 221 tests was tabulated in **Supplementary Table 1** and in the corresponding figure legend. In 55 tests, observations were pooled across animals with no animal-level random effect or cluster-robust correction; those tests were flagged in **Supplementary Table 1** and their p-values were reported descriptively, not as animal-level inferences. The principal claims were also tested at animal or recording level: the neuron-level coupling between hourglass encoding and infraslow entrainment (n = 1,627 units) holds in 12 of 14 mice (Wilcoxon signed-rank p = 0.005; **Extended Data Fig. 9g**), and NREM-to-REM phase-gating takes the mouse as the unit (**Extended Data Fig. 10g**).

#### Per-test audit

**Supplementary Table 1** lists all 221 tests, grouped by the 50 main-figure and Extended Data panels that report them, giving for each the panel, test, unit of inference, n, statistic, effect size and 95% confidence interval, p-value, correction and the source sheet of the deposit.

#### Multiple comparisons

Corrections were applied within pre-specified families; corrected values are reported. Benjamini-Hochberg (BH) correction at q = 0.05 was applied within four p-value families for single-unit state classification pooled across all units (one omnibus, three pairwise; **Fig. 1g,h**); across the nine high-level structures for the firing-rate contrasts (**Fig. 1i**); across the 36 regions tested for P-wave theta-phase gating (**Fig. 6j**); across the 32 regions tested for hourglass encoding (**Fig. 8e,f**); and across the 24 system × metric projection-phenotype tests, 3 of which survive at q = 0.05 (**Fig. 7b–h**). Post-hoc tests following Kruskal-Wallis or Friedman omnibuses were Bonferroni-corrected (**Extended Data Fig. 7a,g**). Uncorrected tests are exploratory and described as nominal associations.

#### Effect sizes and confidence intervals

An effect size accompanies every p-value where one is defined; tests without one are flagged in **Supplementary Table 1**. They are Spearman’s rho, the signed log_2_ fold-change, Kendall’s W, the MRL, and the standardized fixed-effect slope β with 95% confidence interval for every linear mixed model with an estimable standard error and for the meta-analysis, which also reports I^2^ and Cochran’s Q for between-region heterogeneity. Intervals are Wald, or bootstrap percentile where resampling was used (1,000 recording-level iterations, transition analyses; 2,000 animal-level, meta-analysis).

#### Data exclusions

All criteria are quantitative and specified above: automated cluster quality control, no manual curation; ≥10 GOOD units from ≥2 recordings for region inclusion; ≥5 4-s epochs per state for firing-rate comparisons; ≥10 quality-passing neurons, ≥30 epochs per state and ≥1 REM episode for decoder eligibility; and no wake bout >120 s within an hourglass cycle. No data was excluded on any other basis.

#### Replication

Recordings were acquired in 62 penetrations from 22 mice, in 4-h daily sessions over up to four days per animal. Every cohort-level effect derives from multiple animals; the per-animal distributions for the principal claims appear in **Extended Data Figs. 9a, 9g and 10g**.

### AI use in data analysis

Analysis scripts were generated and optimized using Claude Opus (version 4.5 to 5) and Gemini 3 Pro. All AI-generated code was reviewed, validated, and revised by the authors, who confirmed accuracy and reproducibility of results.

## Acknowledgements

This work was supported by the Medical Research Council (MR/V033964/1 and MR/Y004051/1 to S.S.), the European Union’s Horizon 2020 (H2020-ICT, DEEPER, 101016787 to S.S.) and the Rodeki Foundation.

## Author contributions

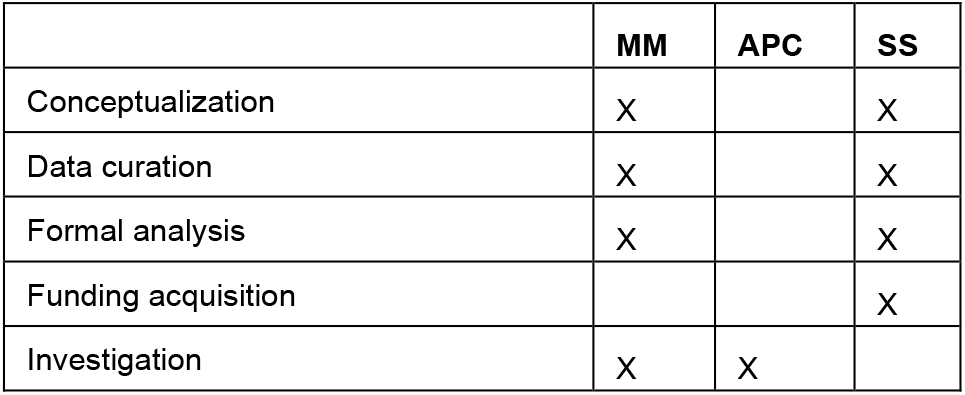
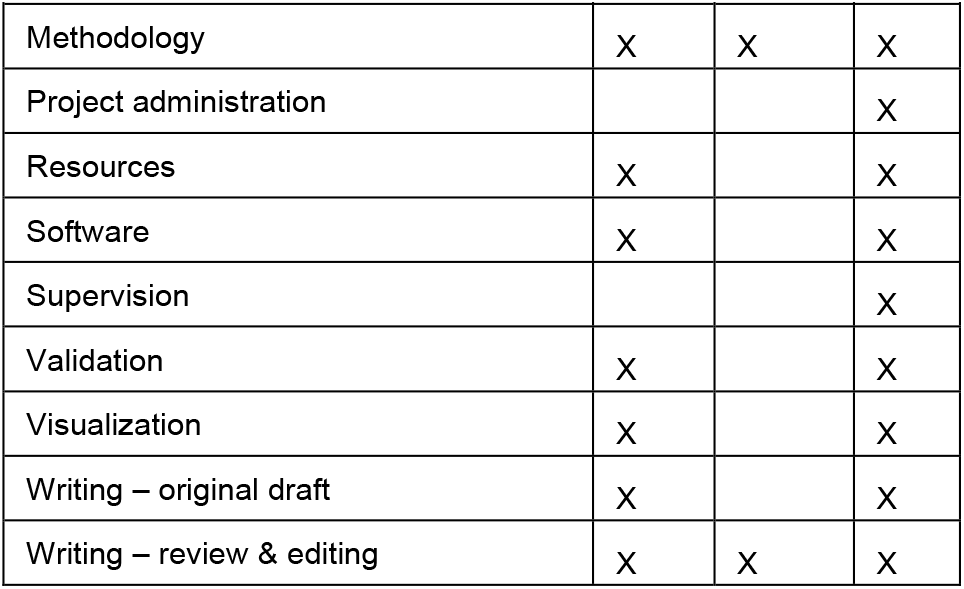

## Extended Data Figures

**Extended Data Figure 1.**
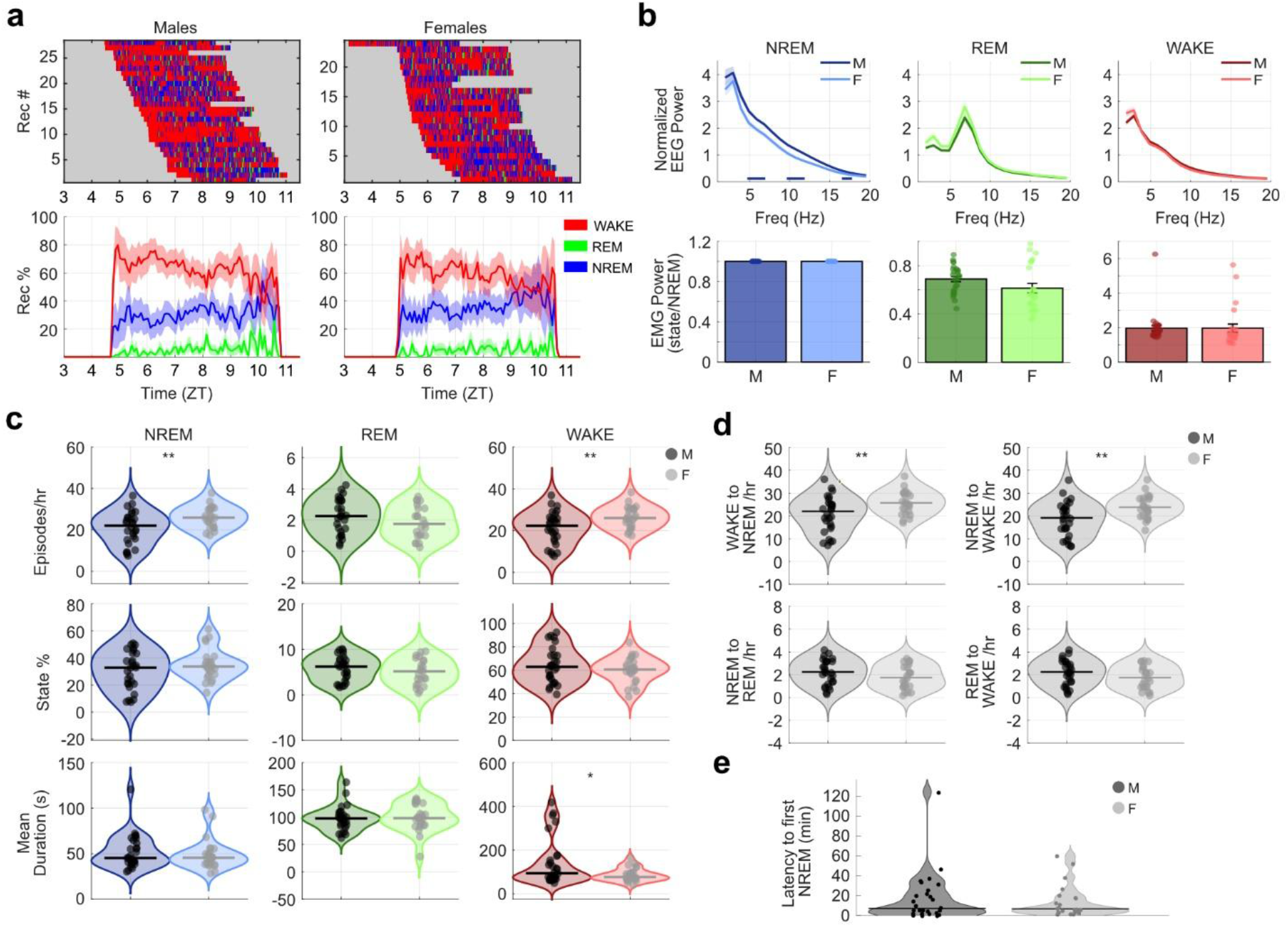
Sleep architecture across datasets. (**a**) Sleep architecture across recordings, split between males (left) and females (right). Top: individual recording hypnograms, bottom: sleep state proportion in recordings across time. (**b**) Average normalized power spectrum density (PSD; top) and normalized EMG power (bottom) for each sleep state, split between males and females. The lines at the bottom of the NREM PSD plot indicate significant differences in power. (**c**) Number of episodes per hour, state percentage, and mean duration of sleep states, split between males and females. (**d**) Number of state transitions per hour for males and females. (**e**) Latency to first NREM episode for males and females. Significance throughout: * p < 0.05, ** p < 0.01.

**Extended Data Figure 2.**
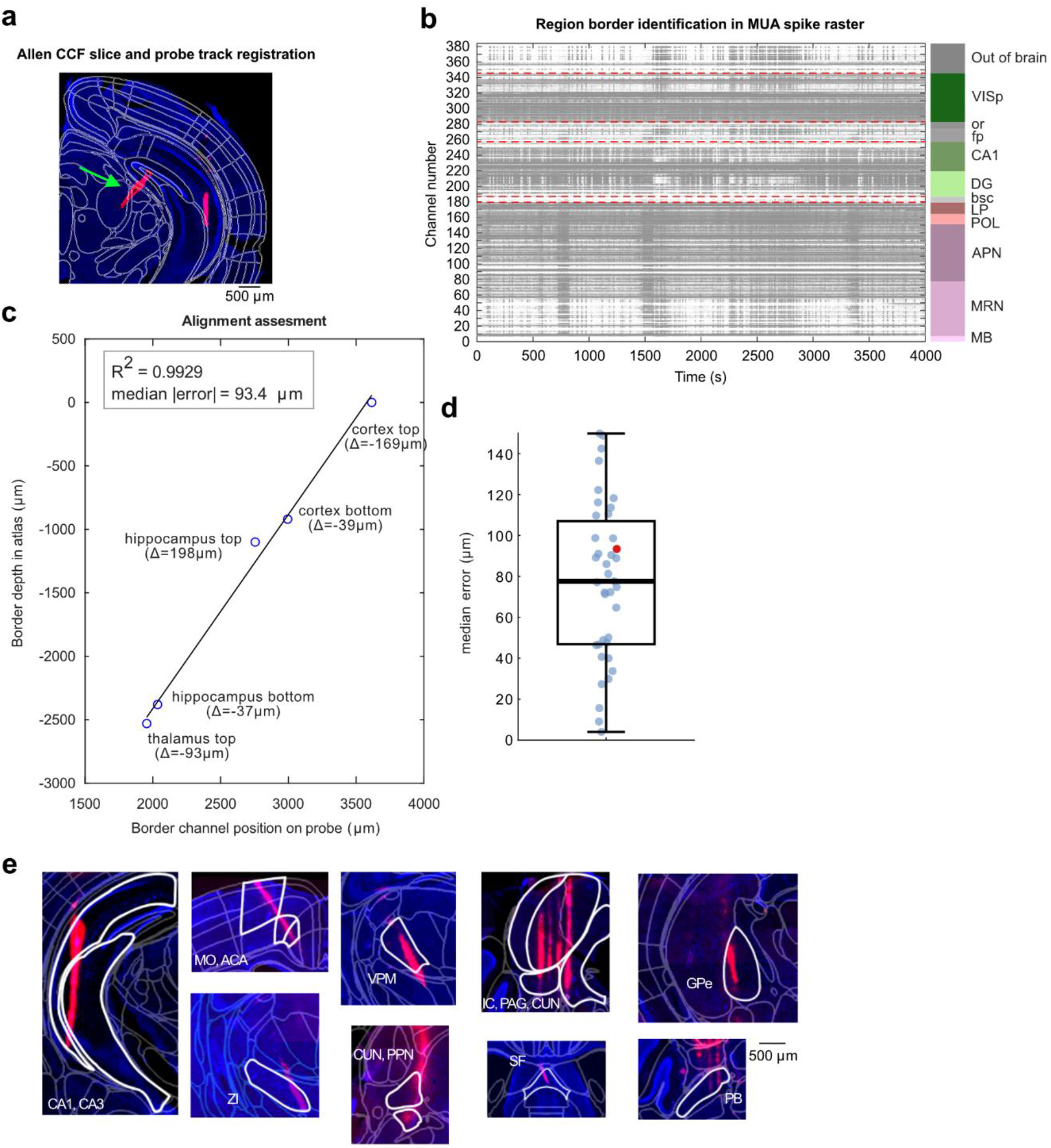
Probe trajectory registration. (**a**) Histological slices were run through SHARP-Track, for atlas alignment and probe track labelling. (**b**) The output of SHARP-Track was a list of brain regions and their estimated borders in µm (right). In parallel, MUA spike raster was plotted across channels (left), and specific region borders were identified from changes in activity (red lines). In this example, 5 borders were found: cortex top (VISp onset), cortex bottom (VISp L6 end and white matter onset), hippocampus top (white matter end and CA1 onset), hippocampus bottom (DG end and white matter onset), and thalamus top (white matter end and LP onset). (**c**) To cross-validate the border locations, we fit a linear regression (with intercept) between border channel positions on the probe and the border Allen CCF atlas locations, and assessed its fit using a leave-one-out cross-validation approach. Prediction errors (residuals) across all landmarks were summarized using the median absolute deviation (for the example dataset, the error is 93.4 µm). (**d**) Boxplot showing median error across datasets (the error of the example dataset is shown as red dot). (**e**) Examples of DiI tracks aligned with the Allen CCF atlas showing recording locations across major brain structures (hippocampus, cortex, hypothalamus, thalamus, midbrain, striatum, pallidum, pons).

**Extended Data Figure 3.**
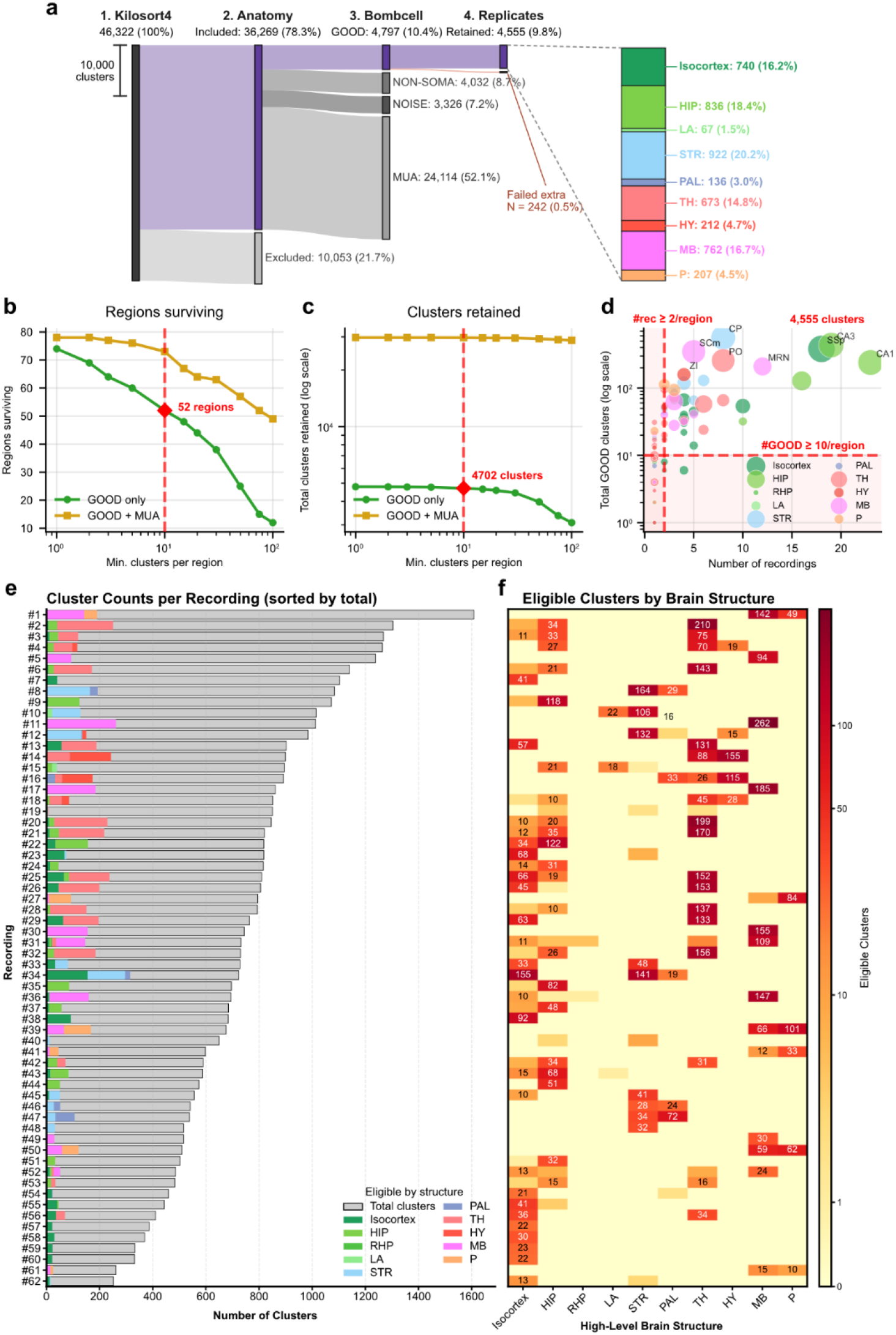
Spike sorting and single-unit quality control. (**a**) Sankey diagram of the cluster-filtering pipeline, retaining 4,555 single units from 46,322 Kilosort 4 clusters (9.8%). Clusters passed sequentially through anatomical inclusion (36,269 assigned to a valid in-brain Allen Common Coordinate Framework (CCF) region; 10,053 excluded as fiber-tract, ventricular or off-target), Bombcell classification (GOOD 4,797, non-somatic 4,032, noise 3,326, multi-unit activity (MUA) 24,114) and a region-validity criterion that removed 242 GOOD clusters lying in border regions. The right-hand column shows the retained units by high-level CCF structure (isocortex, hippocampal formation, lateral amygdala, striatum, pallidum, thalamus, hypothalamus, midbrain and pons). (**b–d**) Selection of the region-level inclusion criterion. In **b** and **c**, the minimum number of GOOD clusters required per brain region is swept (x-axis, log scale); green, GOOD units only; gold, GOOD plus MUA. (**b**) Number of CCF regions surviving. (**c**) Total clusters retained (y-axis, log scale). (**d**) Sampling versus yield: for each region, total GOOD clusters (y-axis, log scale) against the number of recordings sampling it, with bubble area proportional to the region’s total cluster count (all classes) and color denoting high-level structure. Red dashed lines and shading mark the chosen criterion (≥ 10 GOOD units pooled across ≥ 2 recordings), which retained 52 regions and 4,702 GOOD units from the full cluster pool; the anatomically restricted set in a gives the final 4,555 units. (**e,f**) Per-recording single-unit yield across the 62 recordings, ordered by total cluster count. (**e**) Stacked horizontal bars: gray, total Kilosort4 clusters per recording; colored segments, retained single units broken down by high-level CCF structure (colors as in **a**). (**f**) Matrix of retained single units per recording (rows, ordered as in **e**) and high-level structure (columns); shading encodes count (log scale) and counts ≥ 10 are annotated. Across recordings the median retained yield was 63.5 units (range 0–288; 4,555 units in total).

**Extended Data Figure 4.**
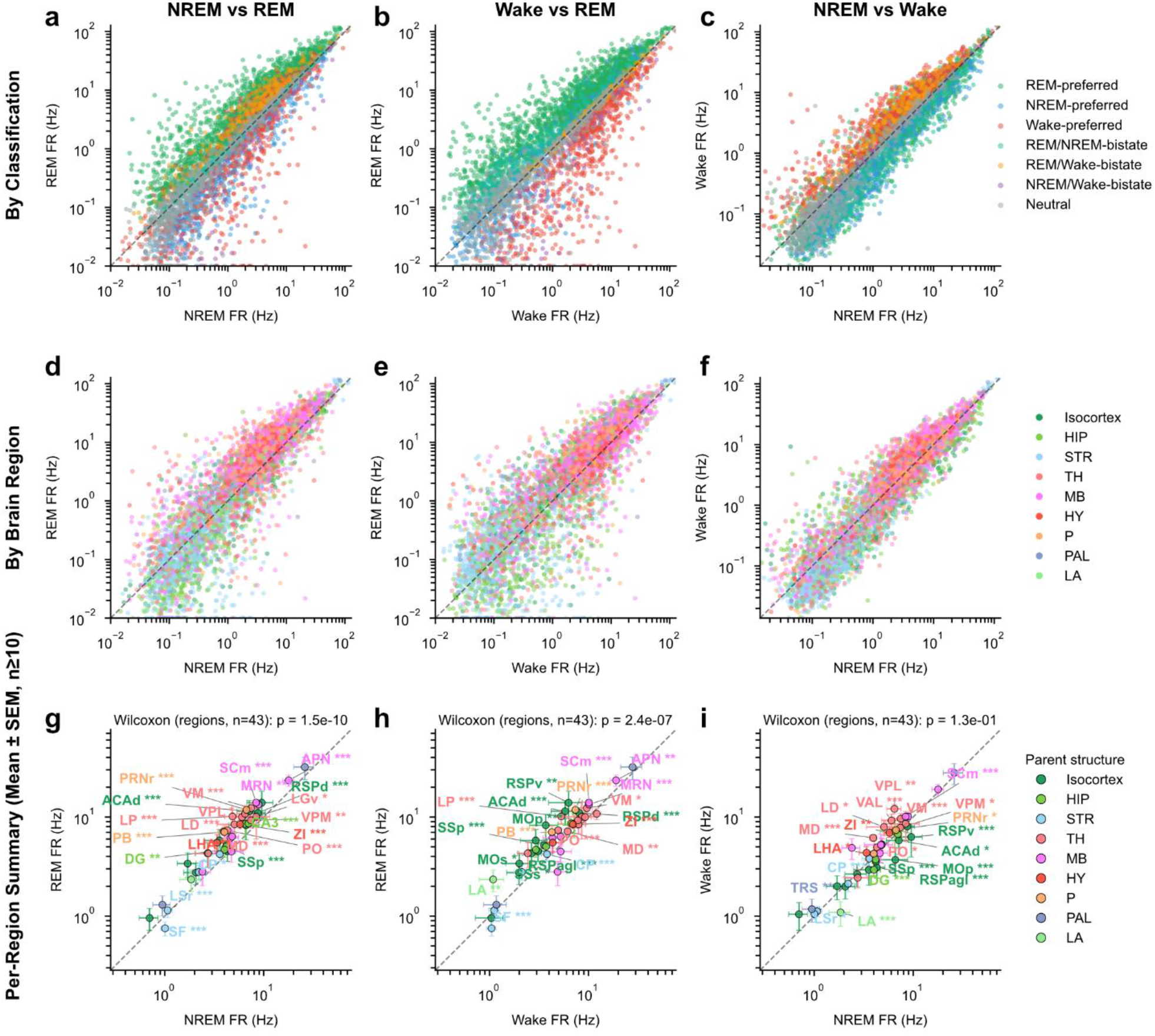
Single-unit firing rates compared across vigilance states. Pairwise comparison of mean single-unit firing rates between vigilance states. Each point is one single unit, plotted at its mean firing rate (Hz, log scale) in the two states indicated – computed as the mean, across that state’s non-overlapping 4-s epochs, of the per-epoch rate; the dashed line marks unity. Analyses include the 4,441 quality-passing units with ≥ 5 epochs in every state (from 59 of the 62 recordings and 43 Allen CCF regions). Columns compare NREM versus REM (REM on the y-axis), wake versus REM (REM on the y-axis) and NREM versus wake (wake on the y-axis). (**a-c**) Units colored by state-preference classification (FDR-corrected). Each unit was assigned to one of seven categories – REM-preferred (n = 1,848), Wake-preferred (750), NREM-preferred (571), REM/NREM-bistate (318), REM/Wake-bistate (250), NREM/Wake-bistate (104) or Neutral (600) – using a permutation framework (10,000 permutations of epoch state-labels): an omnibus ANOVA-style F-test for any state modulation, followed for modulated units by two-tailed pairwise tests of the difference in mean rate for each state pair, with the direction set by the sign of the log_2_ fold-change. A unit was single-state preferred when it fired significantly higher than both other states, bistate when two states fired similarly (|log_2_ fold-change| < 0.5, i.e. within ∼1.41-fold) and both exceeded the third, and Neutral otherwise. All p-values were Benjamini–Hochberg FDR-corrected (α = 0.05) before classification. (**d-f**) The same units colored by high-level CCF structure – isocortex, hippocampal formation (HIP), striatum (STR), thalamus (TH), midbrain (MB), hypothalamus (HY), pons (P), pallidum (PAL) and lateral amygdala (LA). (**g-i**) Per-region summary. Each point is one CCF region with ≥ 10 units (43 regions), plotted at its across-unit mean firing rate ± s.e.m. in the two states, colored by parent structure and labeled with its acronym. Asterisks beside each region denote a paired Wilcoxon signed-rank test of log_10_ firing rate across that region’s units between the two states, Benjamini– Hochberg FDR-corrected across regions. The heading gives a global one-sample Wilcoxon signed-rank test on the per-region log-fold differences (n = 43 regions): NREM versus REM p = 1.5 × 10^−10^, wake versus REM p = 2.4 × 10^−7^ and NREM versus wake p = 1.3 × 10^−1^ (not significant) – region-mean rates were systematically higher in REM than in NREM or wake, but similar between NREM and wake. Significance throughout: *p < 0.05, **p < 0.01, ***p < 0.001; n.s., not significant. Related to Fig. 1i.

**Extended Data Figure 5.**
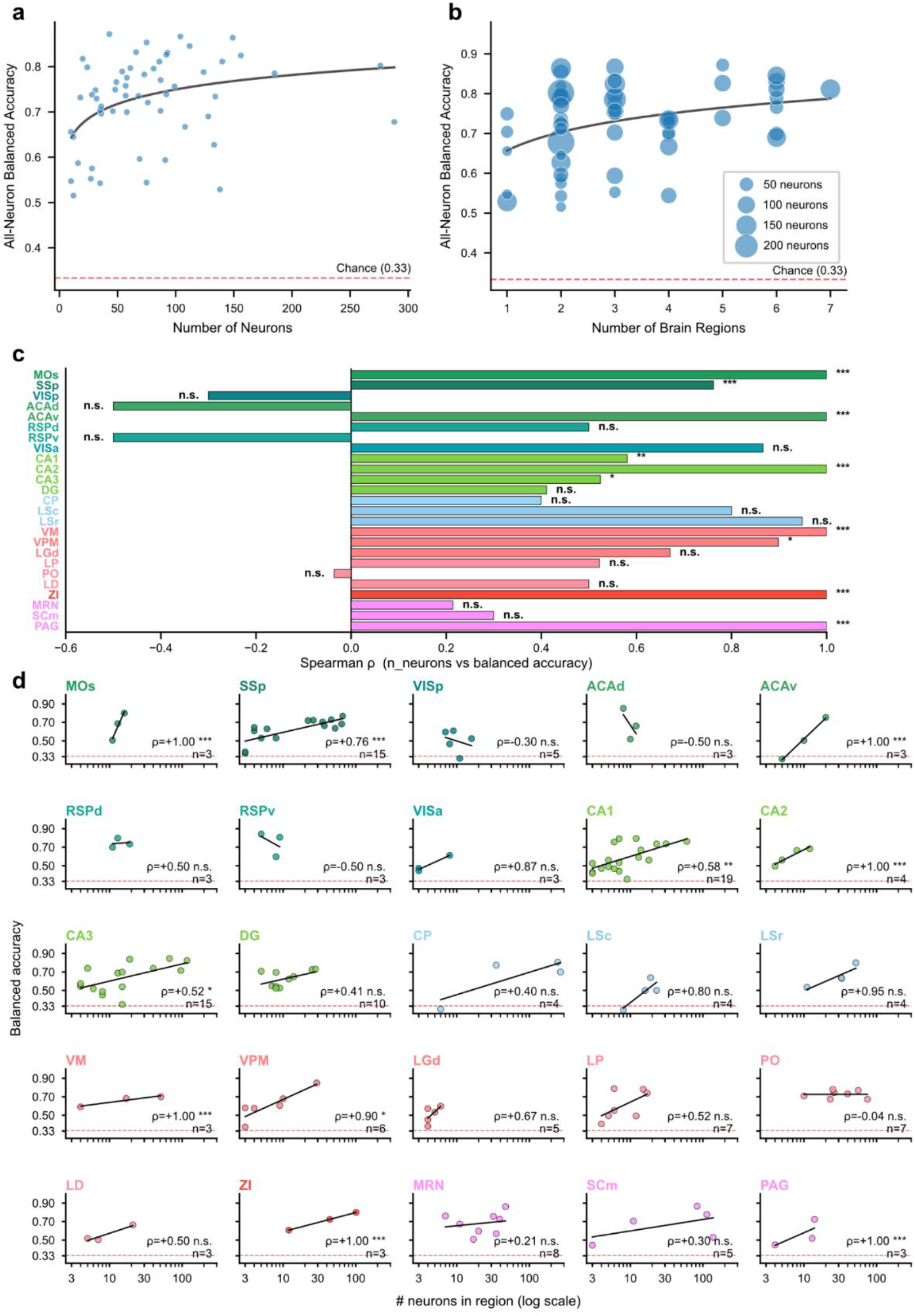
Scaling of vigilance-state decoding with population size and regional coverage. (**a**) All-neuron balanced accuracy—from a decoder using every simultaneously recorded unit—versus the number of units, one point per recording (n = 57). Accuracy increased modestly with population size (Spearman ρ = 0.34, p = 0.010); solid line, logarithmic fit. (**b**) All-neuron balanced accuracy versus the number of brain regions decodable in isolation in that recording (individual Allen CCF regions with ≥ 3 units); marker area scales with unit count (legend, 50– 200 units). Accuracy increased with regional coverage (Spearman ρ = 0.30, p = 0.024, n = 57); solid line, logarithmic fit. (**c,d**) Within-region scaling from decoders trained on the units of one brain region at a time, restricted to the 25 regions sampled in ≥ 3 recordings. (**c**) Per-region Spearman correlation between the number of units in a region and its balanced accuracy across recordings; bars are colored by high-level CCF structure (isocortex, hippocampus, striatum, thalamus, hypothalamus and midbrain) and share the region order of **d**. (**d**) The underlying relationships (balanced accuracy versus unit count, log x-axis; one point per recording; black line, linear fit; red dashed, chance), each annotated with Spearman ρ, its significance and the number of recordings (n). Positive unit-count scaling was significant in ten regions (MOs, SSp, ACAv, CA1, CA2, CA3, VM, VPM, ZI and PAG), most robustly in the densely sampled somatosensory cortex (SSp, n = 15) and hippocampus (CA1, n = 19; CA3, n = 15); coefficients from regions with only three recordings should be interpreted cautiously. Significance throughout: *p < 0.05, **p < 0.01, ***p < 0.001; n.s., not significant.

**Extended Data Figure 6.**
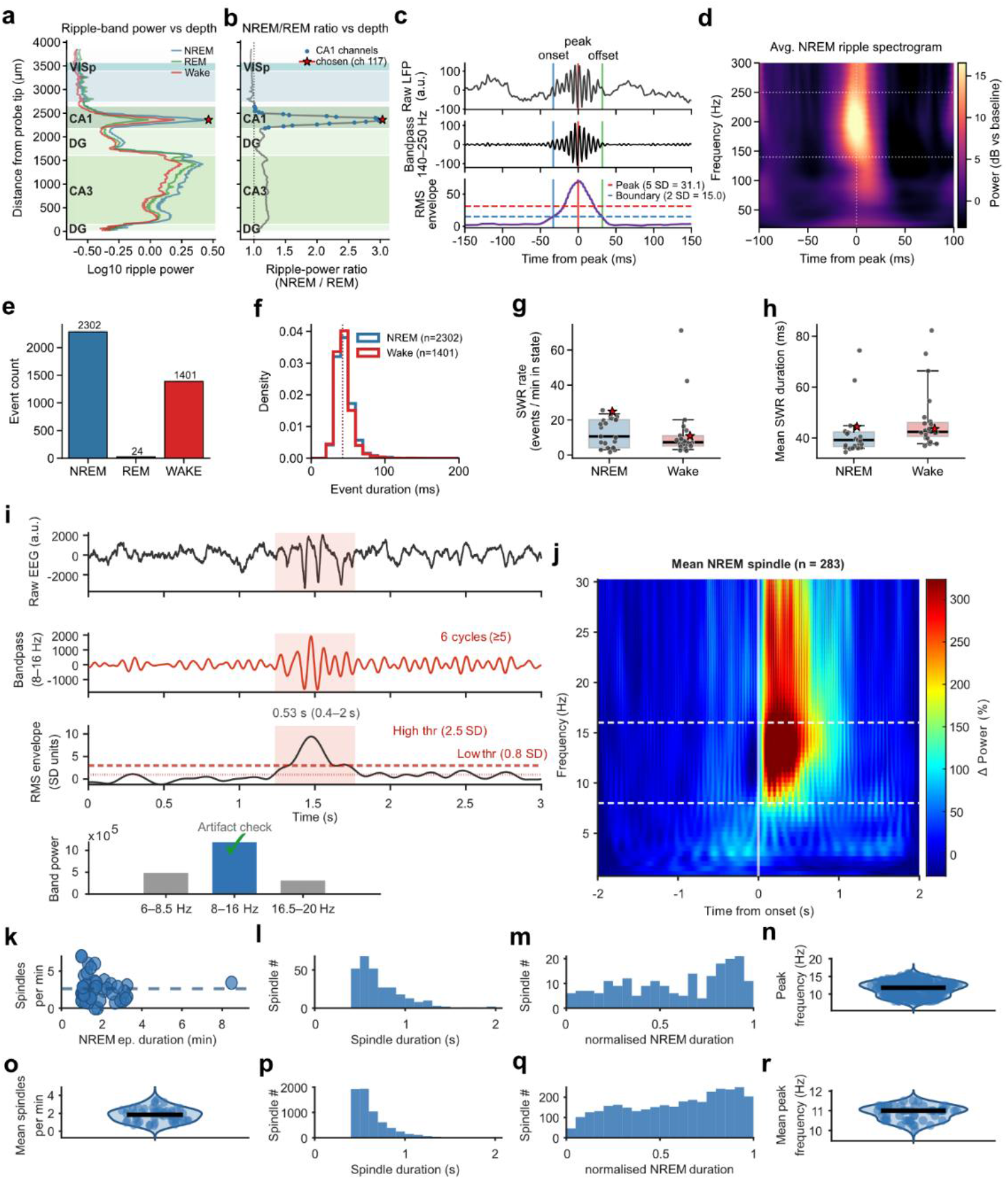
Hippocampal sharp-wave ripples (SWRs) and sleep spindle detection. (**a**) Ripple-band (140–250 Hz) power along the probe, separately for NREM (blue), REM (green) and Wake (red); Allen CCF regions are shaded (visual cortex, VISp; hippocampal CA1, dentate gyrus, DG, and CA3). The detection channel (red star) is the CA1 channel with the highest NREM/REM ripple-power ratio. (**b**) NREM/REM ripple-power ratio versus depth (dotted line, ratio = 1); CA1 channels, blue dots; chosen channel, red star. (**c**) Dual-threshold detection illustrated on one NREM ripple, aligned to its envelope peak: top, raw local field potential (LFP) in arbitrary units. Middle, the 140–250 Hz zero-phase band-pass trace (third-order Butterworth, filtfilt). Bottom, its 20 ms root-mean-square (RMS) envelope. Events were detected where the envelope exceeded a peak threshold (mean + 5 SD; red dashed) and delimited where it fell below a boundary threshold (mean + 2 SD; blue dashed), both estimated from NREM epochs only (here 31.1 and 15.0 a.u.); onset (blue), peak (red) and offset (green) are marked, and the event duration is annotated (65 ms). Accepted events had durations of 20–200 ms; overlapping events were merged. (**d**) Peak-aligned average Morlet-wavelet spectrogram of the NREM ripples from this recording, in decibels relative to the per-frequency pre-event baseline; spectral power concentrates within the 140–250 Hz band (white dotted lines) at the peak (t = 0). (**e**) Event counts by state for the example recording, split into accepted events (NREM, n = 2,302; Wake, n = 1,401; REM, n = 24). (**f**) Event-duration distributions by state for the same recording (NREM and Wake; REM omitted, n = 24 events). (**g,h**) Cohort summary across the 23 recordings (boxes, median and interquartile range across recordings; gray points, individual recordings; red star, the example recording): SWR rate per minute of each state (**g**) and mean SWR duration (**h**), for NREM versus Wake. SWRs occurred at a higher rate during NREM than Wake. (**i**) Spindle detection pipeline: top – raw EEG signal, middle top – bandpass-filtered EEG, middle bottom – signal envelope after Morlet wavelet transform, averaging and smoothing, bottom – band power check for artifact rejection. Detection threshold for an event in the signal envelope was set to 2.5 SD above mean (“high threshold”; thick dotted horizontal line), while onset/offset threshold was set to 0.8 SD above mean (“low threshold”, thin dotted horizontal line). Spindle events had to last between 0.4 and 2 sec, have between 5 and 30 cycles in the bandpass-filtered signal, and power in the spindle band had to exceed the power in the adjacent bands. (**j**) Spectrogram of spindle events in one example recording. (**k-n**) Event quantification for the example recording. (**k**) Spindle rate relationship with NREM episode duration. Horizontal dotted line represents the mean spindle rate for this recording. (**l**) Histogram of spindle duration. (**m**) Spindle number across the (normalized) duration of NREM episode. (**n**) Peak power frequency for spindles detected in the recording. (**o**-**r**) Event quantification across recordings. (**o**) Mean spindle rate across datasets. (**p**) Histogram of spindle duration (pooled events across datasets). (**q**) Spindle number across the (normalized) duration of NREM episode (pooled events across datasets). (**r**) Mean peak power frequency for spindles across datasets.

**Extended Data Figure 7.**
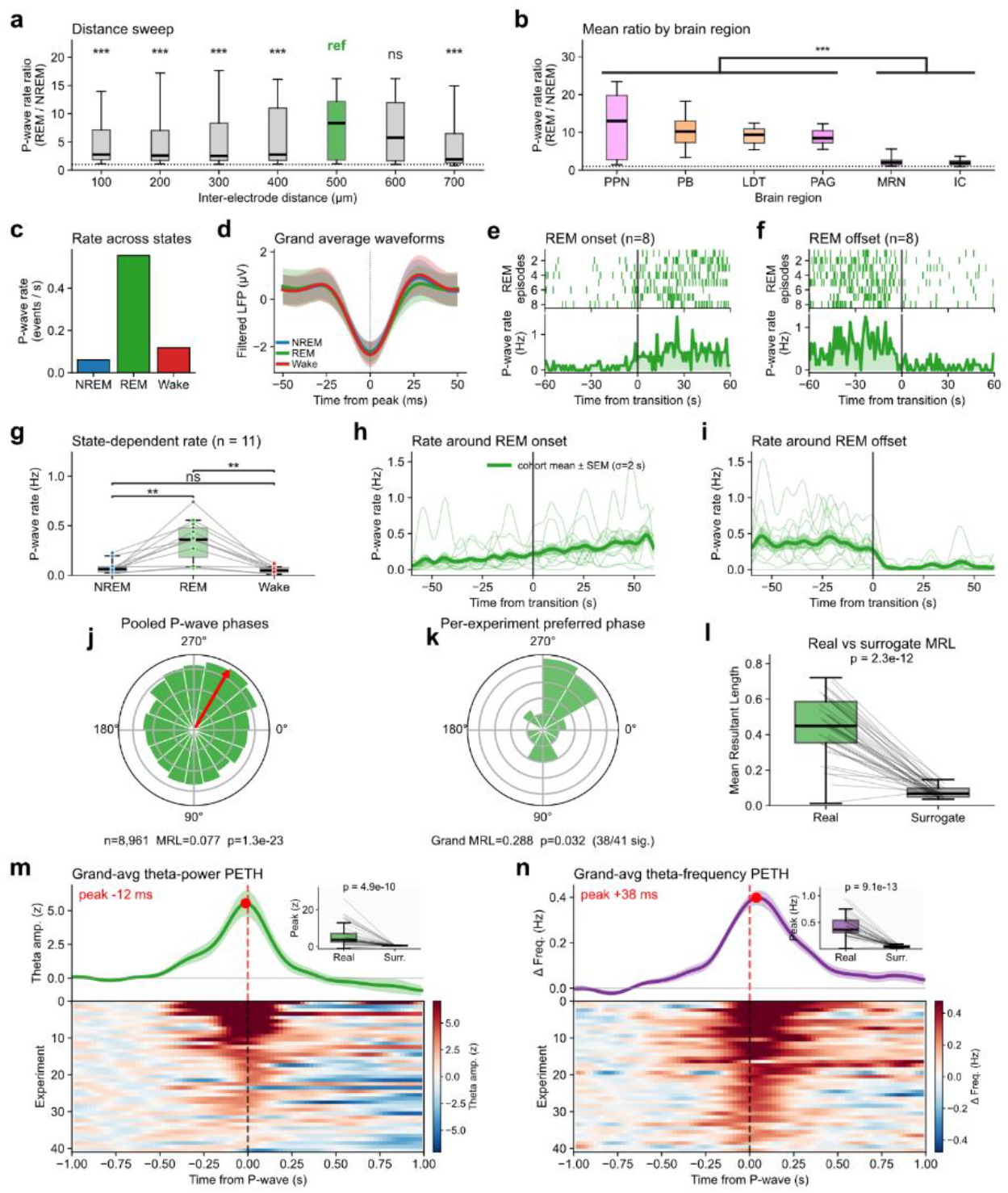
P-wave detection from brainstem Neuropixels recordings and functional coupling of P-waves with cortical theta rhythms during REM sleep. **(a-i**) Detection and validation of P-waves from brainstem Neuropixels recordings. (**a**) Selection of the bipolar interelectrode distance. P-wave REM/NREM rate ratio per electrode pair (state rate, events per second within the state) versus vertical inter-electrode separation (100–700 µm, 100-µm bins; n = 9,837 pairs from 3 recordings in the channel-pair optimization sweep). The 500-µm bin (green, “ref”) was selected as the production distance. Kruskal–Wallis H = 126.7, p = 6.5 × 10^−25^; asterisks, post-hoc two-sided Mann–Whitney U versus 500 µm with Bonferroni correction (***p < 0.001; ns, not significant). (**b**) P-wave REM/NREM rate ratio by brainstem region for the same pairs (each pair contributes to both of its regions), ordered by descending median: PPN (pedunculopontine nucleus), PB (parabrachial nucleus), LDT (latero-dorsal tegmental nucleus), PAG (periaqueductal grey), MRN (midbrain reticular nucleus) and IC (inferior colliculus); box fills follow Allen CCF colors. Regions split into a HIGH (PPN, PB, LDT, PAG) and a LOW (MRN, IC) group (one-sided Mann–Whitney U, Cliff’s δ = 0.92; Kruskal–Wallis across regions H = 12,231.5, p < 10^−300^). The pontine nuclei PPN, PB and LDT were used as the preferred detection sites. (**c-f**) Worked example (PPN–PPN bipolar pair, 495 µm apart; 1,530 detected events). (**c**) Mean P-wave rate per state (events per second): NREM 0.06, REM 0.56, Wake 0.12. (**d**) Grand-average detected waveforms (mean ± s.d. of the filtered LFP) by state, ±50 ms around the negative peak (n = 416 NREM, 382 REM and 732 Wake events). (**e,f**) P-wave times aligned to REM onset (**e**) and REM offset (**f**) across the recording’s 8 REM episodes: per-episode raster (top) and population rate (Hz; ±60 s, 1-s bins). The rate rises at REM onset and falls at REM offset. (**g-i**) Cohort validation across the n = 11 recordings with Neuropixels-derived P-wave detections. (**g**) State-dependent P-wave rate (events per second; box and connected points, one line per recording): REM exceeds NREM and Wake (Friedman χ2 = 18.7, *p* = 8.58 × 10^−5^; post-hoc Wilcoxon signed-rank with Bonferroni correction, \*\**p* < 0.01 for REM versus NREM and REM versus Wake; NREM versus Wake, not significant). (**h,i**) P-wave rate around REM onset (**h**) and REM offset (**i**) (±60 s, 1-s bins, Gaussian-smoothed, σ = 2 s); thin lines, individual recordings; thick line and shaded band, cohort mean ± s.e.m. (**j-n**) phase-locking of pontine P-waves to cortical theta, with transient increases in theta amplitude and frequency around the P-wave. (**j**) Distribution of theta phase at REM P-wave times, pooled across all recordings (n = 8,961 P-waves); the pooled distribution was significantly non-uniform, with a low MRL of 0.077 (Rayleigh p = 1.4 × 10^−23^; red arrow, population mean vector). (**k**) Preferred theta phase of each recording; preferred phases were themselves consistent across recordings (grand MRL = 0.288, Rayleigh p = 0.032), and 38 of 41 recordings were individually phase-locked (Rayleigh p < 0.05). (**l**) Theta phase-locking strength (MRL) for real P-wave times versus matched-N surrogates (random REM times equal in number to each recording’s P-waves); real MRL greatly exceeded surrogate (one-sided paired Wilcoxon signed-rank p = 2.3 × 10^−12^; gray lines, individual recordings). (**m**) Top, grand-average peri-P-wave theta amplitude (mean ± s.e.m.; ±1 s, 25-ms bins; z-scored to a baseline 1.0–0.5 s before the P-wave), which peaked 12 ms before the P-wave (red dashed line, P-wave onset; inset, real versus surrogate peak amplitude, one-sided paired Wilcoxon signed-rank p = 4.9 × 10^−10^). Bottom, per-recording theta-amplitude peri-event time histograms (PETHs; rows, one per recording, sorted by peak amplitude; color, z-scored amplitude). (**n**) Top, grand-average peri-P-wave change in instantaneous theta frequency (Δ frequency relative to baseline; ±1 s, 25-ms bins), which peaked 38 ms after the P-wave (inset, real versus surrogate peak change, one-sided paired Wilcoxon signed-rank p = 9.1 × 10^−13^). Bottom, per-recording theta-frequency PETHs (rows, one per recording, sorted independently by peak frequency change; color, Δ frequency).

**Extended Data Figure 8.**
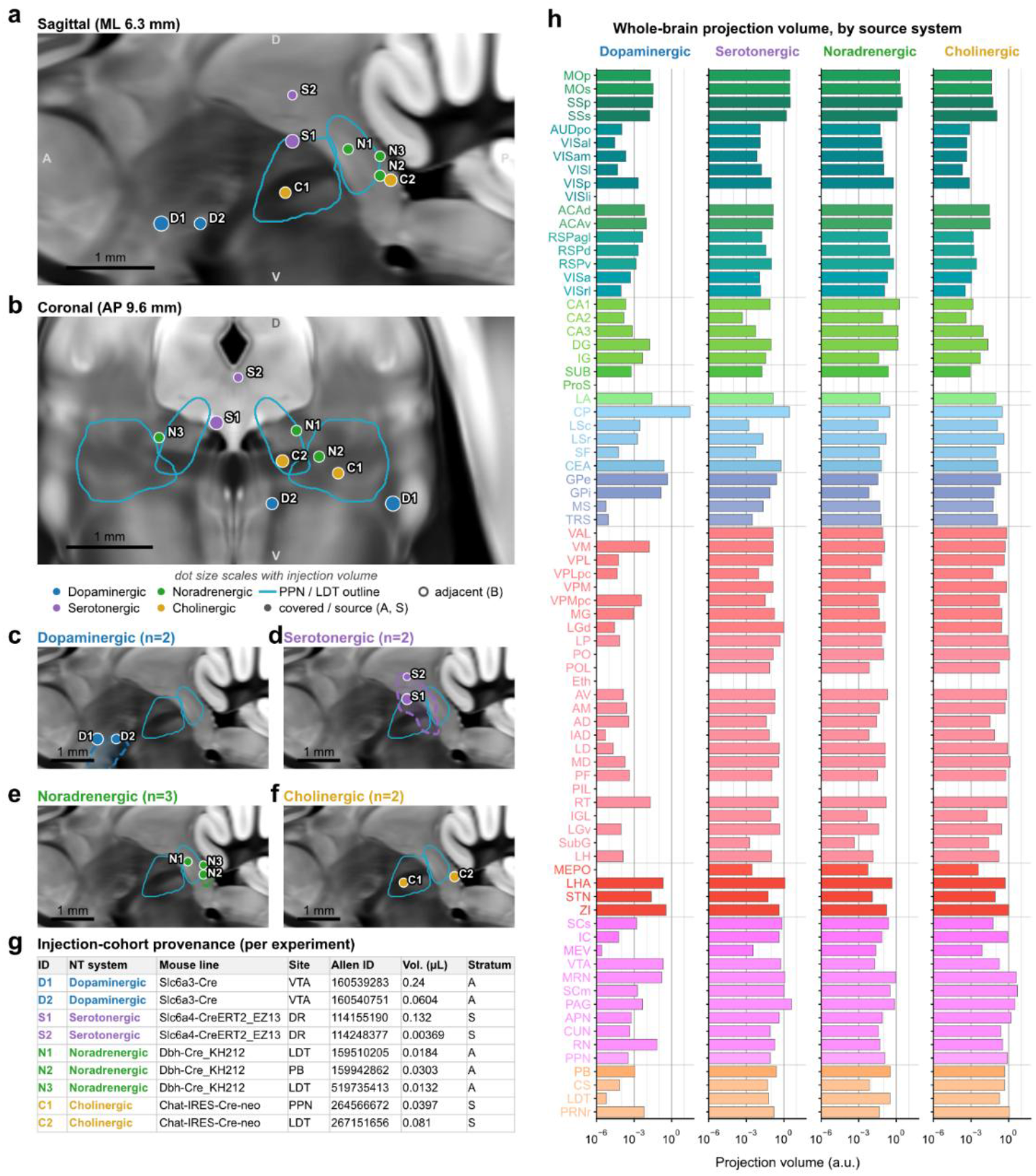
Whole-brain projection volumes across Allen CCF regions for the neuromodulatory systems. Anterograde tracer experiments from the Allen Mouse Brain Connectivity Atlas (n = 9 experiments) used to quantify the cell-type-specific projection densities of the four ascending neuromodulatory systems analyzed in Fig. 7: dopaminergic (DA; Slc6a3-Cre), serotonergic (5-HT; Slc6a4-CreERT2), noradrenergic (NA; Dbh-Cre) and cholinergic (ACh; Chat-IRES-Cre). (**a,b**) All nine injection sites plotted on a representative sagittal (**a**; ML 6.3 mm) and coronal (**b**; AP 9.6 mm) section of the Allen Common Coordinate Framework (CCFv3) average template (25 µm), colored by neuromodulator system (colors as in Fig. 7); the pedunculopontine (PPN) and laterodorsal tegmental (LDT) nuclei — the mesopontine cholinergic source territory — are outlined in teal. The section planes are the cohort-median injection coordinates in Allen CCF space (not stereotaxic). Marker area scales with injection volume (area proportional to the square root of injected tracer volume); scale bar, 1 mm. (**c–f**) The same experiments shown separately by system on the sagittal section, each with that system’s biological source nucleus outlined and lightly shaded in the system color: dopaminergic, ventral tegmental area (VTA; **c**); serotonergic, dorsal raphe (DR; **d**); noradrenergic, locus coeruleus (LC; **e**); cholinergic, PPN + LDT (**f**). Nuclear outlines are Allen CCF annotation boundaries drawn as thin-slab maximum projections through the plane; panel headings give the number of experiments per system (DA, n = 2; 5-HT, n = 2; NA, n = 3; ACh, n = 2). Filled dots denote stratum A (the injection envelope covers the target nucleus) or stratum S (a biology-defined source region). Each injection carries a system-prefixed identifier (D, dopaminergic; S, serotonergic; N, noradrenergic; C, cholinergic) shown beside every dot and in the table; the noradrenergic experiments were selected as locus-coeruleus-covered injections, so their primary Allen structure labels (LDT or parabrachial nucleus, PB) are the injection centroids listed in **g**. (**g**) Per-experiment provenance: neuromodulator system, mouse (Cre) line, primary injection structure, Allen Mouse Brain Connectivity Atlas experiment ID, injection volume (µL) and stratum. Because nuclear outlines are slab maximum projections and injection sites are projected onto the shared section, out-of-plane (third-axis) position is collapsed. (**h**) Region-wise anterograde projection volume for four tracer source systems, shown as horizontal bars on a base-10 logarithmic axis (projection volume in arbitrary units, a.u.). Data are anterograde Cre-driver tracer experiments from the Allen Mouse Brain Connectivity Atlas, queried programmatically (AllenSDK). For each target region, projection volume is the normalized projection volume (projection signal within the target divided by the injection-site volume) averaged across that system’s qualifying experiments; finer Allen structures were aggregated (summed) into their parent recorded region, giving 78 CCF regions. Experiments were grouped by injection-source nucleus and mouse (Cre) line: dopaminergic (DA; Slc6a3-Cre; ventral tegmental area and substantia nigra; n = 2), serotonergic (5-HT; Slc6a4-CreERT2; dorsal raphe; n = 2), noradrenergic (NA; Dbh-Cre; locus coeruleus; n = 3), and cholinergic (ACh; Chat-IRES-Cre; pedunculopontine and laterodorsal tegmental nuclei, PPN/LDT; n = 2); Related to Fig. 7a.

**Extended Data Figure 9.**
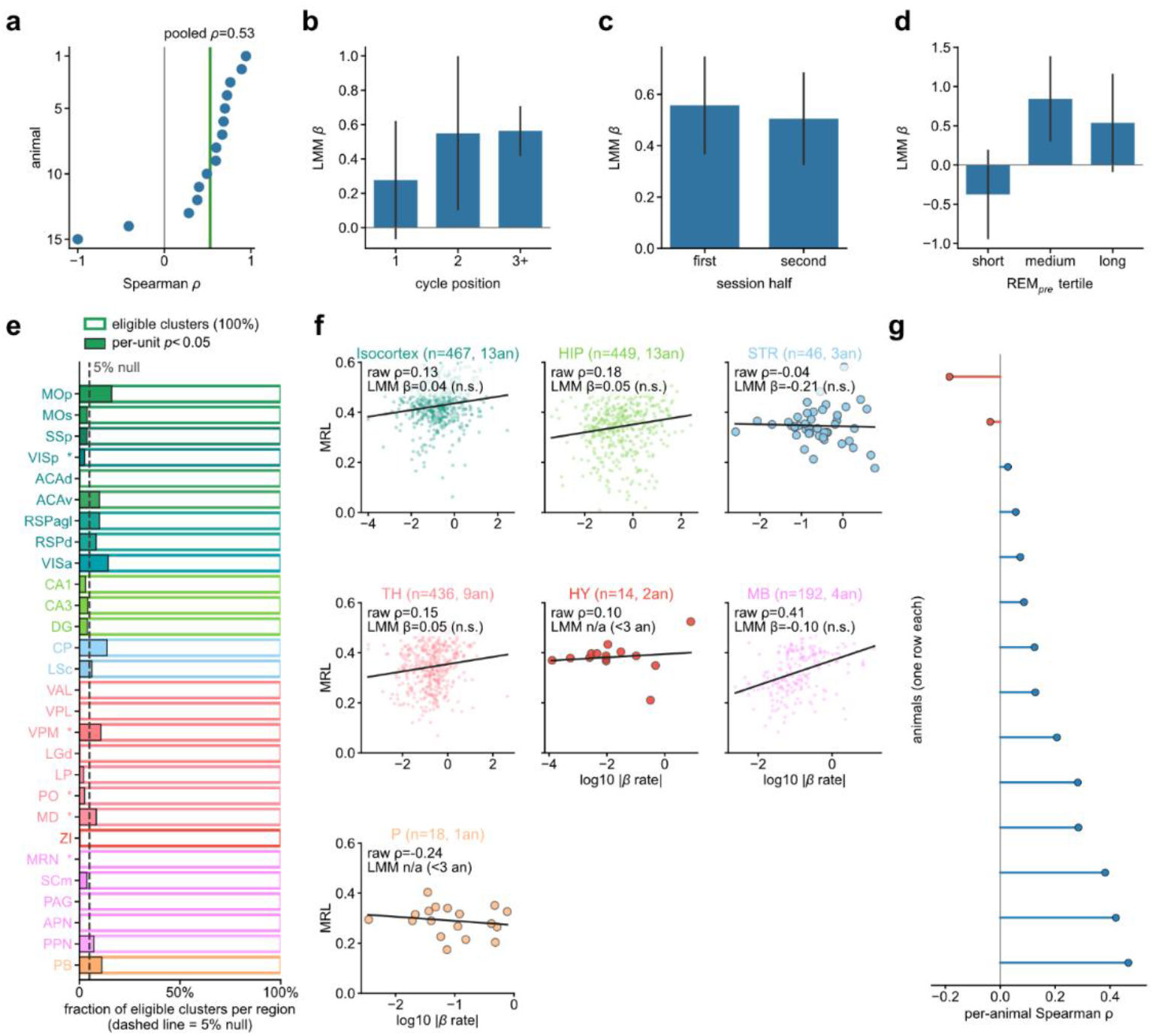
Robustness of the behavioral hourglass relationship, single-unit permutation testing of hourglass discharge-encoding and its coupling to infraslow spindle-band activity. (**a–d**) Robustness of the behavioral relationship (158 inter-REM cycles from 19 mice; cycles interrupted by a wake bout longer than 120 s were excluded). Throughout, the dependent variable is log-transformed NREM accumulation (log |N|) and the predictor is log-transformed preceding-REM duration (log REM_pre_). (**a**) Per-animal Spearman correlation between REM_pre_ duration and subsequent NREM duration; one point per mouse, restricted to 15 of 19 mice with at least three cycles. The correlation was positive in 13 of 15 mice; the green line marks the pooled correlation across all 158 cycles (ρ = 0.53, p = 9.8 × 10^−13^). (**b–d**) Fixed-effect slope of log |N| on log REM_pre_ (LMM β) estimated within strata from a linear mixed model with a per-animal random intercept (ordinary-least-squares fit where the mixed model did not converge); error bars, 95% confidence intervals. The relationship was robust across (**b**) cycle position within the recording (first, second and third-or-later inter-REM cycle; β = 0.28, 0.55 and 0.56; n = 40, 33 and 85 cycles) and (**c**) session half (first versus second half of the recording; β = 0.56 and 0.51; n = 68 and 90 cycles). Splitting by (**d**) tertiles of REM_pre_ duration (short, medium and long; β = −0.38, 0.84 and 0.54; n = 55, 50 and 53 cycles) showed that it was carried by the medium and long REM episodes and absent for the shortest ones. (**e**) Per-region single-unit “yield” of the hourglass discharge-encoding test, showing that a small number of individual units reach significance. For every quality-passing single unit, the test asked whether its mean firing rate during the preceding REM episode (REM_pre_) predicted the amount of NREM sleep that subsequently accumulated before the next REM episode (|N|), over and above REM_pre_ duration, using a residualized-rate ordinary-least-squares model evaluated against a within-recording permutation null (500 permutations; uncorrected per-unit p < 0.05). Each horizontal bar is one Allen CCF region with at least ten eligible units (n = 28 regions, 1,609 units). The vertical dashed line at 5% marks the chance (null) expectation. Across the full cohort-wide test (1,629 units with a valid permutation fit, from 14 mice and 26 recordings) only a single unit survived Benjamini– Hochberg false-discovery-rate correction (q < 0.05). Asterisks mark the five plotted regions that were nevertheless significant in the hierarchical, region-level linear mixed-model analysis, including MRN, which reached hierarchical significance although none of its 45 units passed the per-unit test; two further hierarchically significant regions (TRS and LD) had fewer than ten eligible units and are not shown. (**f,g**) Coupling between a unit’s hourglass discharge-encoding and its entrainment to the infraslow spindle-band rhythm, across 1,627 quality-passing single units from 14 mice. For each unit, hourglass discharge-encoding is the magnitude of its discharge-encoding coefficient (|β_rate_|), the ordinary-least-squares coefficient of the unit’s mean firing rate during REM_pre_ on the subsequent |N|, over and above REM_pre_ duration, and infraslow entrainment is the mean resultant length (MRL) of its NREM spike phases relative to the infraslow (0.01–0.05 Hz) envelope of σ/spindle-band (10–15 Hz) EEG power. (**f**) MRL versus log_10_ |β_rate_| for every unit, grouped by Allen CCF high-level structure: isocortex, hippocampal region (HIP), striatum (STR), thalamus (TH), hypothalamus (HY), midbrain (MB) and pons (P). Each panel is annotated with the raw within-structure Spearman ρ and, where a structure contained at least 30 units from at least three mice, the standardized adjusted coefficient (LMM β) from the mixed model z(MRL) ∼ z(log_10_ |β_rate_|) + z(log_10_ firing rate) + (1 | animal), which controls for firing rate and between-animal differences (labelled ‘n/a’ for structures with fewer than three mice). No individual structure was significant. (**g**) Consistency across mice: the per-animal Spearman correlation between |β_rate_| and MRL (mice with at least 20 units) was positive in 12 of 14 mice (two-sided Wilcoxon signed-rank p = 0.005). Pooled across all units the coupling was positive and significant (mixed-model β = 0.094, p = 2.3 × 10^−4^, n = 1,627 units, 14 mice; raw Spearman ρ = 0.145), indicating a weak, distributed coupling rather than one localized to any single structure.

**Extended Data Figure 10.**
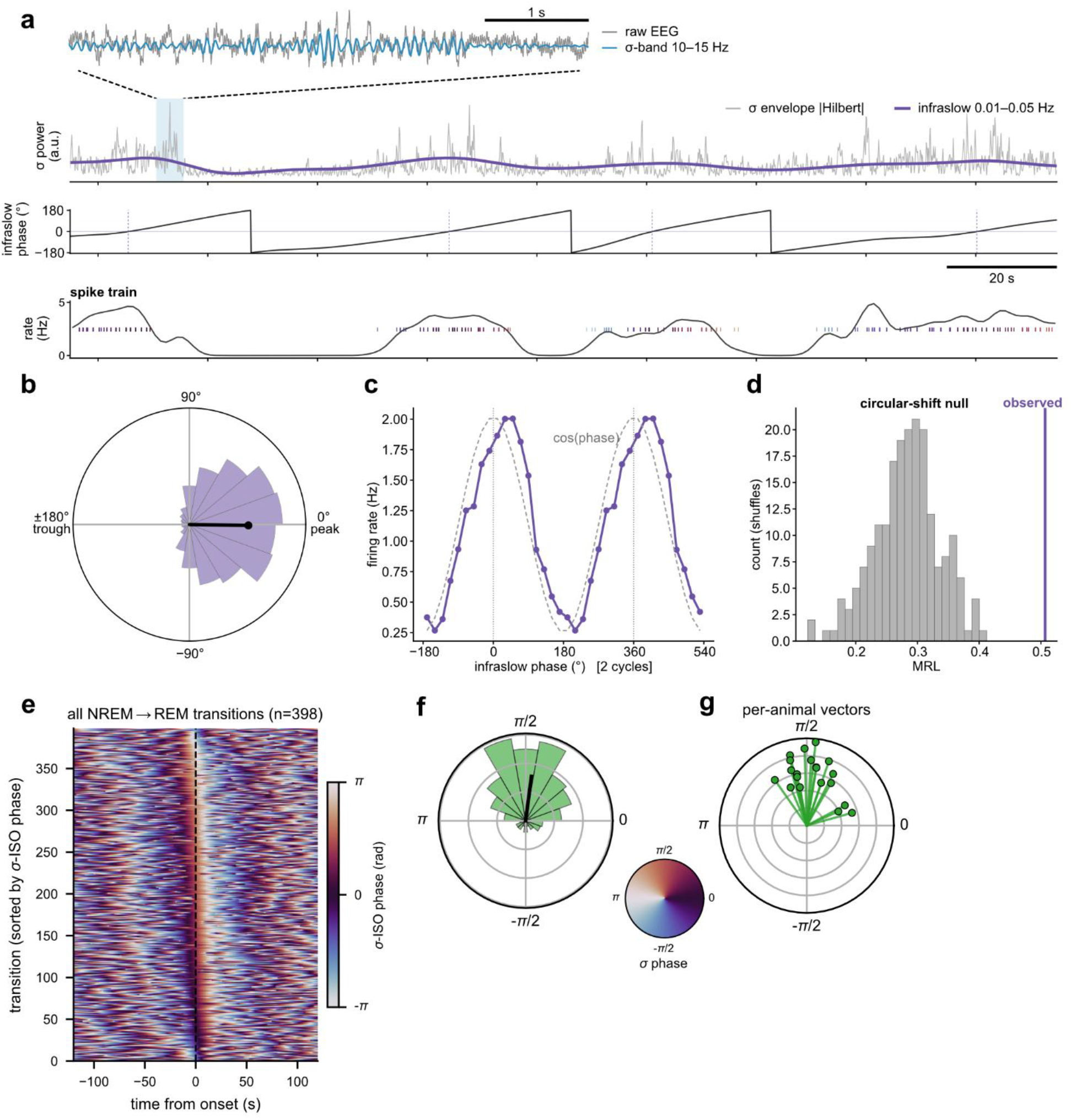
The infraslow oscillation of σ (spindle-band) power entrains neuronal firing and gates NREM-to-REM transitions. (**a–d**) Entrainment of single-unit firing to the sigma-infraslow (σ-ISO) oscillation, illustrated for a representative unit. (**a**) The σ-ISO extraction pipeline applied to this unit: raw EEG and its 10–15 Hz σ-band component (5-s zoom at a σ-power peak); the σ-band amplitude envelope and its 0.01–0.05 Hz infraslow band-pass; the Hilbert phase of the infraslow envelope (0° = σ-power peak); and each NREM spike coloured by the σ-ISO phase assigned at its spike time by circular interpolation (n = 6,681 NREM spikes). (**b**) Distribution of σ-ISO phases across the unit’s NREM spikes with the mean resultant vector (black); mean resultant length (MRL) = 0.51, preferred phase φ = −1° (firing concentrates essentially at the σ-power peak). (**c**) Occupancy-normalized firing rate versus σ-ISO phase (spike count per 20° bin divided by the time spent in that bin during NREM; two cycles shown) with a cosine reference (dashed); firing is maximal near the σ-power peak (0°/360°). (**d**) The observed MRL (0.51; purple line) against a circular-shift null distribution (gray; k = 200 surrogates, each generated by a single random circular time-shift of the spike train by ≥ 60 s and recomputing the MRL), giving a surrogate z-score of 4.3. (**e–g**) The σ-ISO phase gates the timing of NREM-to-REM transitions across the dataset (48 recordings from 22 mice; simultaneously recorded probes collapsed to one series per recording; a NREM-to-REM transition is a NREM epoch immediately followed by a REM epoch). (**e**) σ-ISO phase in a ±120-s window (1-s bins) around every NREM-to-REM transition, one row per transition, rows sorted by the σ-ISO phase at onset (t = 0, dashed line); the 398 of 400 transitions with a complete ±120-s window are shown. A coherent phase band at t = 0 indicates that transitions occur at a preferred σ-ISO phase. (**f**) Pooled distribution of the σ-ISO phase at transition onset (18 bins) with its mean resultant vector (black): transitions are strongly phase-concentrated (Rayleigh R = 0.54, p = 9 × 10^−50^, n = 400 transitions), preferring a phase ∼83° past the σ-power peak. The color wheel (between **f** and **g**) maps σ-ISO phase to color. (**g**) Per-mouse mean σ-ISO onset vectors (one green vector per mouse; vector length is that mouse’s within-mouse MRL; n = 20 mice with ≥ 3 NREM-to-REM transitions). The phase preference is consistent across mice (second-level Rayleigh on the per-mouse mean directions, R = 0.88, p = 1 × 10^−7^); the mouse is the unit of inference, guarding against pseudo-replication.

## Supplementary Information

### Supplementary Files

**Supplementary File. Interactive Swanson flatmap of the mouse brain (Allen CCF).** Browser-based, interactive companion to the static Swanson flatmaps in **Fig. 1e** as a single self-contained HTML file. The complete left-hemisphere Swanson flatmap of the mouse brain is shown in portrait orientation, with every Allen Common Coordinate Framework (CCF) region drawn as a filled polygon and colored by its Allen CCF ontology color, following the high-level-structure palette used throughout the paper (isocortex, hippocampal formation, lateral amygdala, striatum, pallidum, thalamus, hypothalamus, midbrain and pons; cf. **Fig. 1f**). Hovering the cursor over any region reveals its Allen CCF acronym and full anatomical name (for example, PO — Posterior complex of the thalamus), giving an interactive key to the region abbreviations used throughout the figures. Standard plot controls (zoom, pan, autoscale/reset and download as PNG) are available from the toolbar. Region geometry and Allen CCF colors were obtained with iblatlas (Swanson flatmap polygons and the Allen CCF region ontology).

**Supplementary Video. Brain-wide reorganization of population activity across sleep–wake transitions, shown as synchronized Swanson flatmaps and delay-embedded principal-component trajectories.** Animated summary of peri-transition population dynamics for the cohort (22 head-fixed mice, 62 Neuropixels recordings, 4,555 quality-controlled single units assigned to 46 Allen CCF regions spanning nine high-level brain structures). The four columns show, left to right, the four canonical vigilance-state transitions: Wake → NREM, NREM → Wake, NREM → REM and REM → Wake. Each region’s peri-transition firing-rate profile was baseline z-scored (baseline −60 to −30 s), Gaussian-smoothed (FWHM 2 s) and averaged with pure-state masking, so that only epochs whose scored state matched the expected pre- or post-transition state contributed to each time bin. Bottom row: a Swanson flatmap of the mouse brain in which each recorded region is filled by its instantaneous z-scored firing rate (RdBu color map; red, increase above baseline; blue, decrease; symmetric scale saturated at ± 15 z; regions without coverage in grey), with a shared horizontal color bar giving the z-score versus baseline. Top row: the same regional profiles rendered as delay-embedded principal-component analysis (PCA) trajectories in the PC1–PC2 plane (4 s embedding window, 0.5 s step), in which every region traces a path colored by its high-level CCF structure, carrying a trailing history and a leading marker at the current time; axis labels give the variance explained by each component (PC1, 96.8–99.3% across transitions). The animation advances through peri-transition time from −25 to +45 s relative to the scored transition boundary (t = 0) in 500 ms steps at 10 frames per second (14 s total); a per-panel timestamp and a faint background tint mark the current vigilance state (Wake, red; NREM, blue; REM, green) and switch at the transition. At the two arousal-defining boundaries (Wake ↔ NREM) forebrain and brainstem regions move in opposite directions along this near-one-dimensional axis, whereas at the REM transitions they move in register. Related to **Figs. 3 and 4**.

